# Stabilizing selection of seasonal influenza receptor binding in populations with partial immunity

**DOI:** 10.1101/2020.06.27.175190

**Authors:** James A. Hay, Alvin Junus, Steven Riley, Hsiang-Yu Yuan

## Abstract

Mutations that alter cellular receptor binding of influenza hemagglutinin (HA) have profound effects on immune escape. Despite its high mutation rate, it is not fully understood why human influenza HA displays limited antigenic diversity across circulating viruses. We applied phylogenetic analysis and phylodynamic modeling to understand the evolutionary and epidemiological effects of binding avidity adaptation in humans using net charge as a marker for receptor binding avidity. Using 686 human influenza A/H3N2 HA sequences, we found that HA net charge followed an age-specific pattern. Phylogenetic analysis suggested that many binding variants have reduced fitness. Next, we developed an individual-based disease dynamic model embedded with within-host receptor binding adaptation and immune escape in a population with varied partial immunity. The model showed that mean binding avidity was unable to adapt to values that maximized transmissibility due to competing selective forces between within- and between-host levels. Overall, we demonstrated stabilizing selection of virus binding in a population with increasing partial immunity. These findings have potential implications in understanding the evolutionary mechanisms that determine the intensity of seasonal influenza epidemics.

## Introduction

Seasonal influenza viruses cause recurrent epidemics due to the repeated emergence of novel antigenic variants ([1], [2]). This is a result of antigenic drift, whereby immune-mediated selection acting on epitope regions of surface glycoprotein hemagglutinin (HA) allows mutant viruses to escape host immunity induced by previous infections or vaccinations.([1], [3]). Although amino acids in the HA globular head domain that change binding to antibodies are directly selected by host immunity during influenza infection, other factors, such as viral attachment to host cells, also play an important role in immune-evasion and may drive influenza antigenic evolution through frequent alteration of cellular-receptor binding avidity ([4], [5], [6], [7]). This mechanism was first supported by a serial passaging study of pandemic A/H1N1, in which cellular receptor binding avidity increased in immune mice and decreased in naive mice ([4]). These observations suggest that an increase in cellular receptor binding avidity might improve immune escape within an immune host, whereas a reduction in binding avidity within a naive host might bear the cost of impeding the efficiency of viral replication or shedding during infection. This indicates that cellular-receptor binding is itself a trait under immune selection in mice, where the phenotypes that maximise within-host fitness differ between immunologically naive individuals and immune individuals.

Whether receptor binding avidity adapts in humans as it does in mice remains unclear. There is a lack of population studies demonstrating if and how influenza virus binding avidity adapts in human populations, where transmission occurs between individuals with a wide range of immune states. Challenges exist because unlike in serial passaging experiments using nasal injections, humans are mainly exposed to a limited number of virus particles during natural exposure through aerosol or contact transmission ([8], [9]). Thus, the chance of successful infection will largely depend on different levels of clinical protection from pre-existing immunity ([10], [11], [12]). Different human infection and vaccination histories would result in a diverse immune spectrum from weak to strong in a population ([13]), where transmission between naive, immune and partially immune hosts may depend on and select for different binding avidity phenotypes ([14]).

A consequence of within-host adaptation to heterogeneous immunity in a population is that phenotypic clustering (e.g., receptor binding) of viruses could occur based on the immune status of their hosts: viruses isolated from immunologically similar hosts should be phenotypically more similar than viruses isolated from immunologically dissimilar hosts. Studies have been performed to examine the relationship between phylogenetically distinct viral groups and clinical manifestations or age of the patients but no correlation has been shown ([15], [16]). However, no study has been performed to examine clustering of receptor binding of viral isolates based on age-specific immune status of humans or time of isolation. This leaves a number of outstanding questions: is there evidence of influenza receptor binding adaptation to population immunity in the evolutionary history of human influenza viruses? If such adaptation exists, what is its impact on influenza transmission and evolution?

Given the high mutability of influenza ([17], [18]) and frequent occurrence of novel variants within the host ([19], [20]), influenza viruses are paradoxically limited in their genetic and antigenic diversity at the global scale ([21]). A seminal study discovered that only a few substitutions near the receptor binding site determine much of the major antigenic changes in influenza evolution ([22]). Understanding constraints on influenza antigenic evolution may elucidate the potential impact of antigenic variants on future disease outbreaks. However, little is known about the mechanism by which they are constrained.

To address the above questions, we demonstrate here that binding avidity adaptation is likely present in human populations. Phylogenetic analysis of viral sequences suggested that stabilizing selection exists on cellular receptor binding avidity. We developed an epidemic model embedded with within-host receptor binding adaptation and a population structure with varying degrees of immune status to understand the complex interplay between receptor binding and host immunity and their impact on influenza epidemics.

## Results

### Net charge is a good marker of binding avidity

In order to evaluate whether the adaptation of binding avidity to host immunity can be observed within human influenza infection histories, we first investigated whether changes in the net charge of HA can be used as a molecular marker for binding avidity, as suggested previously ([23] [24]). Because sialic acid receptors on host cells have a negative charge, a positive net charge on viral HA is likely to increase the electrostatic force between the viral HA and its sialic acid receptor ([25]). We collected amino acid substitutions and their corresponding receptor binding avidity changes that were either publicly available from the literature or from a laboratory conducting experiments. The receptor avidities of the substitutions were measured using agglutination assays with pre-treated neuraminidase as receptor-destroying enzymes (RDE) ([26], [4]). All derived single substitutions were grouped according to whether they reduced (negative), maintained (neutral), or increased (positive) the net charge of the viral HA (Figure 1) by counting the number of positively charged amino acids minus negatively charged amino acids.

**Figure 1.**
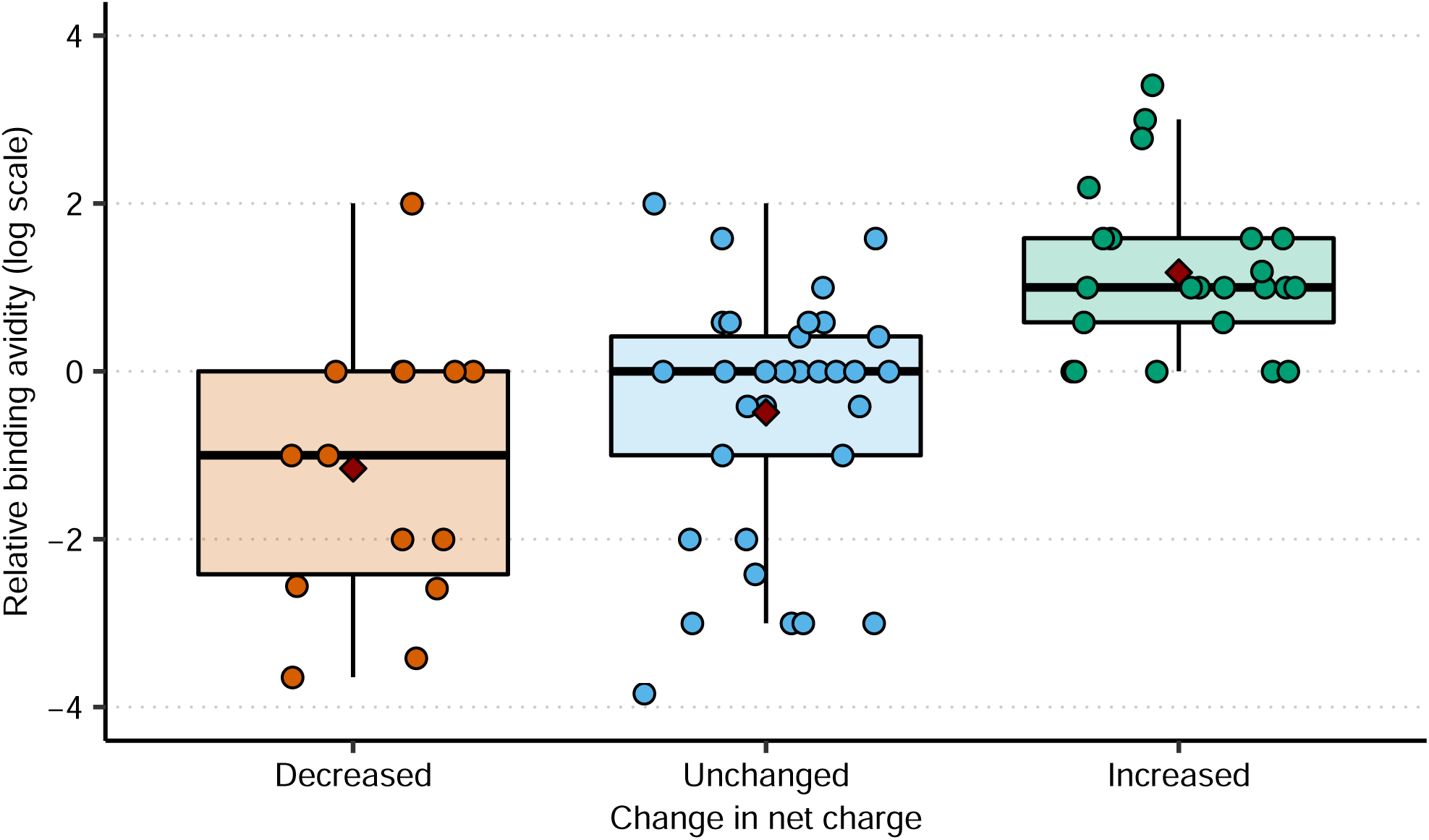
Binding avidity change of single amino acid mutations by net charge. The value is calculated as the log ratio of RDE activity of the mutant to that of the wild type. Amino acid mutations are grouped into 3 categories. *Increase* represents amino acid changes that increase net charge by 1 or 2 units. *Decrease* represents the amino acid changes that decrease net charge by 1 or 2 units. *Unchanged* represents amino acid changes that do not change net charge. Aspartate (D), glutamate (E) were counted as negatively charged amino acids (−1) and Histidine (H), Arginine (R), Lysine (K) were counted as positively charged (+1). Diamonds show mean values.

Substitutions that led to a higher HA net charge enhanced binding avidity compared to mutations that reduced net charge (Table S1). We estimated the odds ratio of an increase in binding avidity from an increase in net charge relative to a decrease or no change to be 11.27 (95% confidence interval (CI): 3.14-48.5; p-value*<*0.001 using Fisher’s exact test), indicating that net charge is a good marker for HA binding avidity. All p-values were calculated using two-sided tests in this study unless specified otherwise. Compared to a reference virus strain PR8 (A/H1N1), there was also a positive correlation between binding avidity and the net charge of HA (Figure S1).

### Net charge correlates to age in humans

To investigate if binding avidity adaptation is present amongst influenza viruses circulating in humans, we obtained 759 influenza A/H3N2 sequences alongside infected persons’ age metadata isolated from 1993-2006 as part of the Influenza Genome Sequencing Project ([27]). The percentage of viruses with a high net charge (defined as an absolute net charge greater than the median value of 18) followed an inverted bell-shaped distribution by age when fitting a smoothing spline (Figure 2). Persons between 20-50 years old (yo) generally had the lowest percentage of virus isolates with high net charge, whereas younger children (¡5 yo) and individuals older than 60 yo exhibited higher percentages of high net charge isolates.

**Figure 2.**
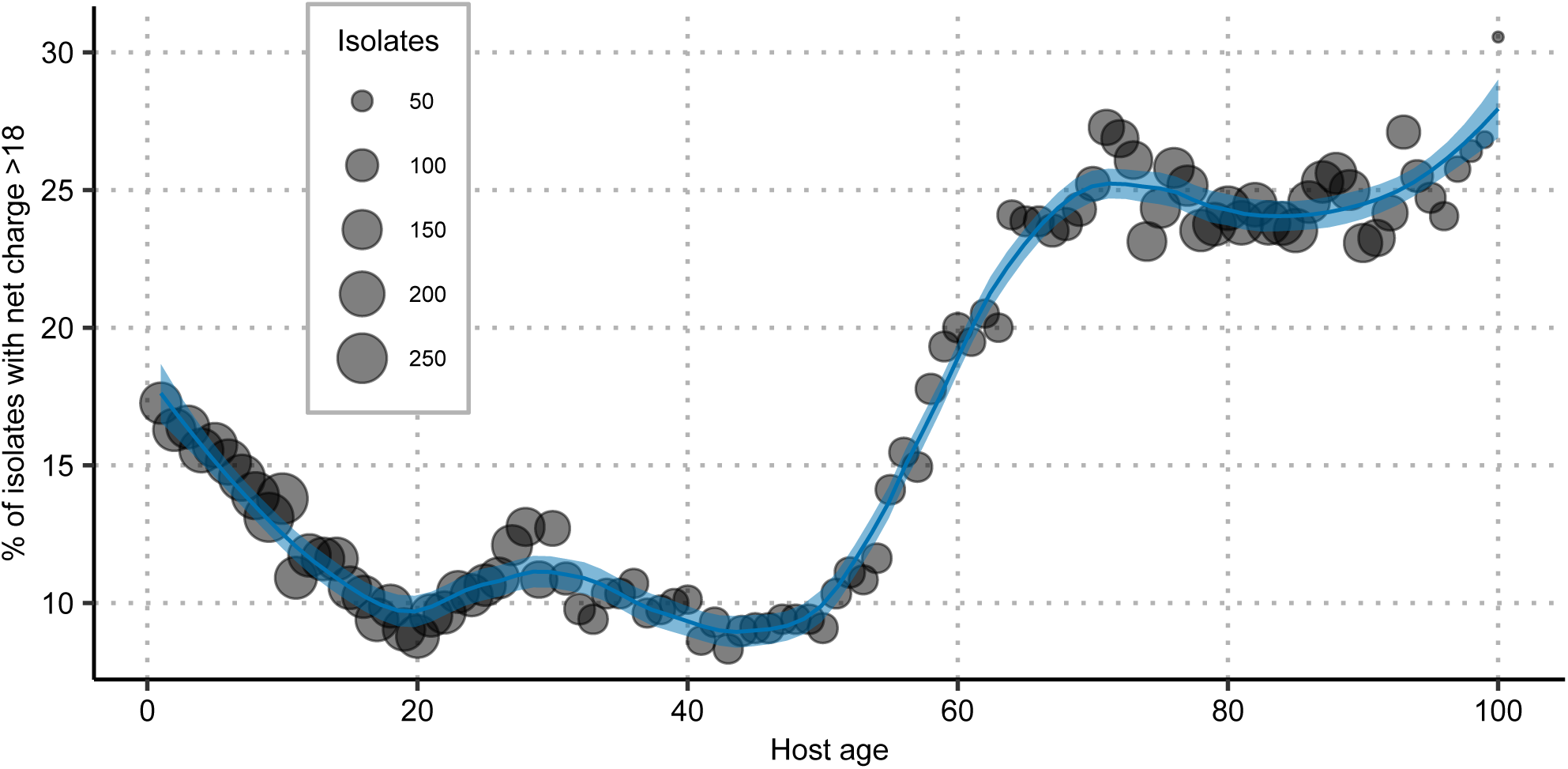
Percentage of high net charge viral isolates by age. Each circle represents the proportion of individuals with high net charge isolates (*>* 18 where median net charge is 17) calculated using a sliding window of size 10 years centered on each year from 1 to 100 yo. The size of circle represents the number of strains in a given sliding window. A smooth curve (blue line) was fitted with the calculated values by LOESS regression using a span value of 0.3. 759 virus sequences and age metadata were obtained from the Influenza Genome Sequencing Project ([27]) and the Influenza Virus Resource Database ([28]).

Both the percentage of viruses with a high net charge and the net charge distribution (Figure S2) of different ages from these human virus isolates showed similar patterns to age specific serological prevalence. Antibody titers against the hemagglutinin head of current influenza A/H3N2 strains are highest in children and the elderly, and lowest in middle aged individuals ([29] [30] [31] [32]). The observed age differences in seroprevalence could be a consequence of original antigenic sin and non-HA targeted immunity leading to lower antibody titers in adults relative to children, and more lifetime exposures with higher vaccination rates in the elderly. These data support the binding avidity hypothesis that a higher binding avidity would be selected for by higher levels of host immunity in a human population as a result of adaptation for immune escape ([4]).

### Phylogenetic analysis of A/H3N2

To further investigate whether binding avidity alterations occurred frequently among transmission events in A/H3N2 infection histories, we used 686 (a subset of the full 759) viral sequences isolated from North America (including sequences from the Influenza Genome Sequencing Project) and reconstructed the phylogenetic tree using BEAST ([33]). The ancestral states of the nucleotides in internal nodes were also inferred during Bayesian reconstruction and were then transformed to amino acid sequences. While we observed changes in net charge between different clades, we did not observe frequent sequential alterations within clades (Figure 3A). After we grouped strains by the number of glycosylation binding sites (#NGS) (which could possibly reduce binding avidity by shielding cellular receptors), net charge of the main trunk was maintained at stable, intermediate values, and changed slowly over time.

**Figure 3.**
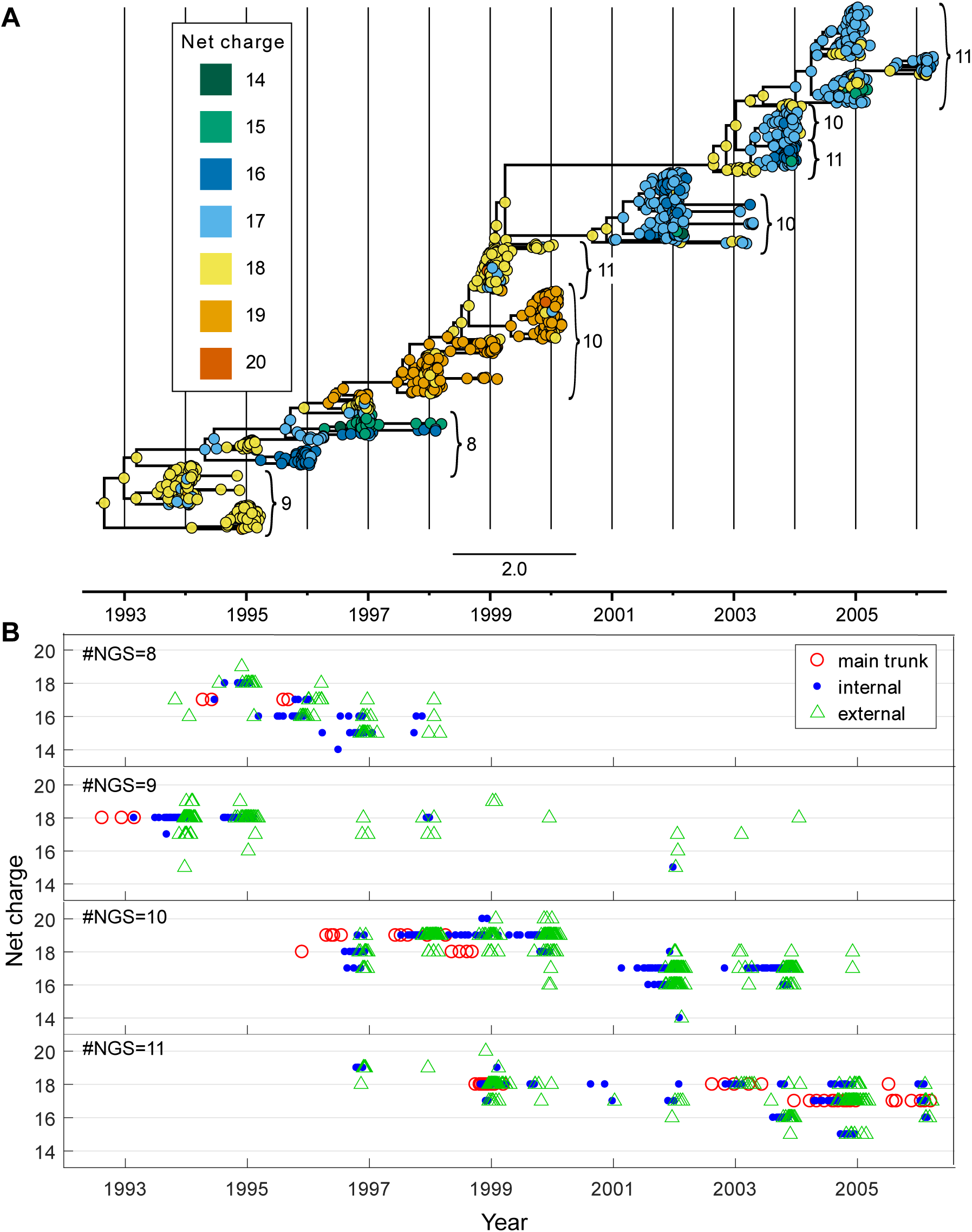
Phylogenetic analysis of cellular receptor binding avidity. (A) The maximum clade credibility (MCC) tree for the A/H3N2 viral isolates from New York State, with HA net charge values mapped onto all inferred internal nodes and all observed external nodes. Numbers next to the bracketed clades denote the #*NGS* group. (B) Viral net charge dynamics over time. Each point represents a node from (A) and is colored by whether it is classified as an internal main trunk node (red circle), an internal non-trunk node (blue dot), or an external node (green triangle). Placement along the x-axis corresponds to the inferred or observed time of the node in the phylogeny. Placement along the y-axis corresponds to calculated net charge value. Each of the four subplots shows viral net charge dynamics for a single #*NGS* group.

A lower number of net charge-produced binding mutations were found in internal nodes than external nodes (Figure 3B). Across all internal nodes, 7.7% of the strains showed net charge changes relative to the reconstructed ancestral sequence; however, a higher proportion of external nodes (12.1%) demonstrated changes in net charge (Table S2). Nodes that demonstrated changes in net charge were approximately 40% less frequent in internal than external nodes. Normalizing by total branch length for internal and external branches yielded similar results, with fewer net charge changes occurring per unit time on internal relative to external branches. The odds ratio for a change in net charge for strains at external nodes relative to internal nodes was 1.64 (95% CI: 1.14-2.35; Fisher’s exact test; p-value=0.009; n=1370). Since external nodes are more likely to die out than internal nodes, these data suggest that many strains with HA binding avidity changes (with avidities that are too high or too low) may have reduced fitness in the population, which is a key characteristic of stabilizing selection.

### The impact of binding avidity adaptation on epidemic dynamics

To measure the role of binding avidity in an influenza epidemic, we first evaluated the impact of binding avidity adaptation on epidemic dynamics using an individual-based (also called agent-based) model to compare the scenarios where the binding avidity trait was or was not allowed to undergo within-host adaptation. Within-host binding avidity adaptation was determined by the within-host fitness gradient conditional on the immunity of the host (Figure 4A), such that binding avidity changed towards the value that maximized the within-host reproductive number during an infection. We assumed that rapid evolution and limited intrahost genetic diversification through new mutations was present ([20]), and therefore we only recorded a single binding avidity trait to represent the virus population in an infected individual. Given a contact, the corresponding probabilities of infection at different levels of host immunity were also determined (Figure 4B). Both within-host fitness and the probability of infection reached a maximum at the same virus binding avidity, indicating that binding avidity adaptation increased not only virus reproduction within a host but also transmission probability between hosts. We therefore hypothesized that binding avidity adaptation would increase incidence during an epidemic.

**Figure 4.**
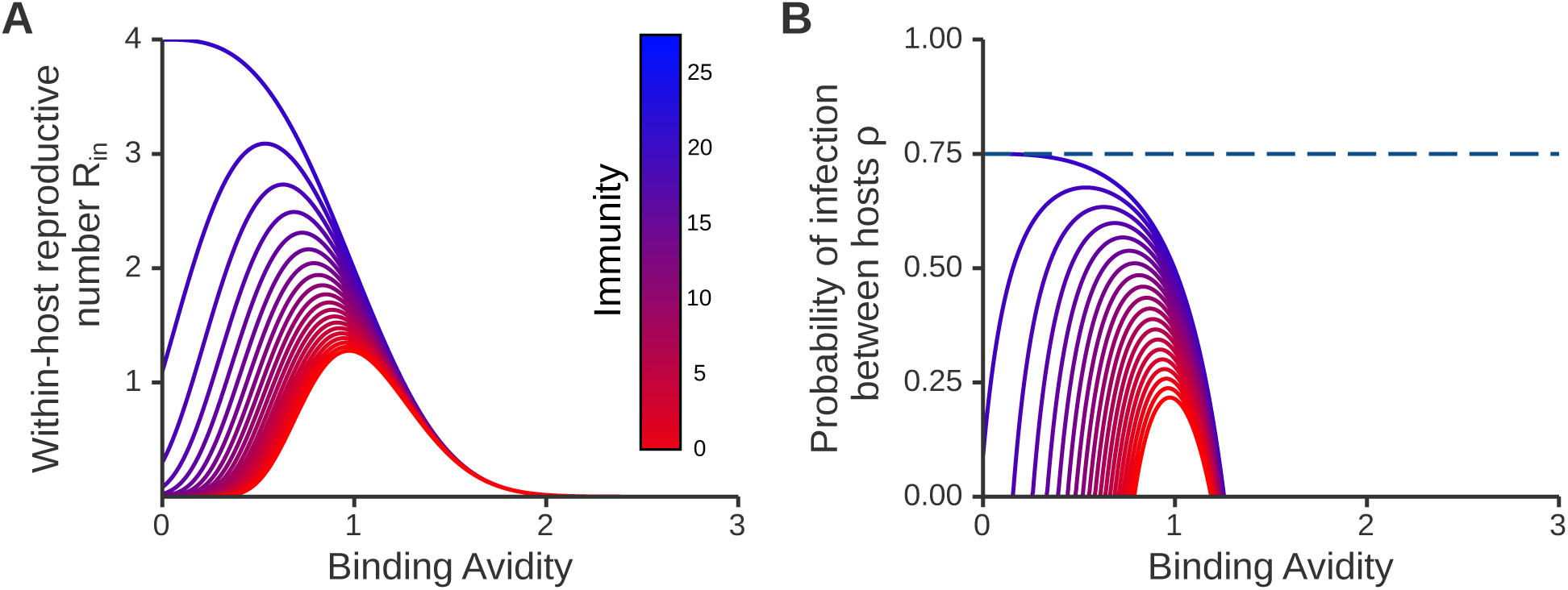
Within-host reproductive number and the probability of infection. (A) Within-host fitness represented by the expected within-host reproductive number *R*_*in*_. Parameters for within-host immune escape and replication cost were set as *p* = 4, *r* = 70, *a* = 0.7 and *b* = 3, and the effective number of virions produced per replication was *n* = 4 (see Table S4 for parameter descriptions). Blue indicates infection in an immunologically naive host, whereas red indicates a host with stronger immunity. (B) The probability of infection to a susceptible host. Within-host parameters values are the same as in (A) and the effective number of transmitted virions was *σ*=1.

We simulated disease dynamics after introducing an antigenically novel virus into a population with varying degrees of pre-existing partial immunity. Incidence reached a peak approximately 200 days after seeding with an antigenically novel virus when the model allowed binding avidity to undergo within-host adaptation (Figure 5A). Regardless of the assumed binding avidity of the initial seed virus, mean binding avidity across extant viruses quickly adapted to a relatively low value (near 0.45) within the first 30 days of the epidemic, indicating rapid adaptation to the lower initial population immunity. Later, mean binding avidity gradually increased toward a higher value (near 0.70) from 50 to 400 days after seeding (Figure S3). This increase in mean binding avidity correlated with an increase in average population immunity (Figure 5B), indicating adaptation to the gradual accumulation of host immunity during the epidemic. The number of immunologically naive individuals (defined as having no immunity against the infecting virus) involved in transmission dropped as successful infections increased the number of people with partial immunity. Therefore, successful infections later in the epidemic were from viruses with higher binding avidities that were adapted to higher levels of partial immunity.

**Figure 5.**
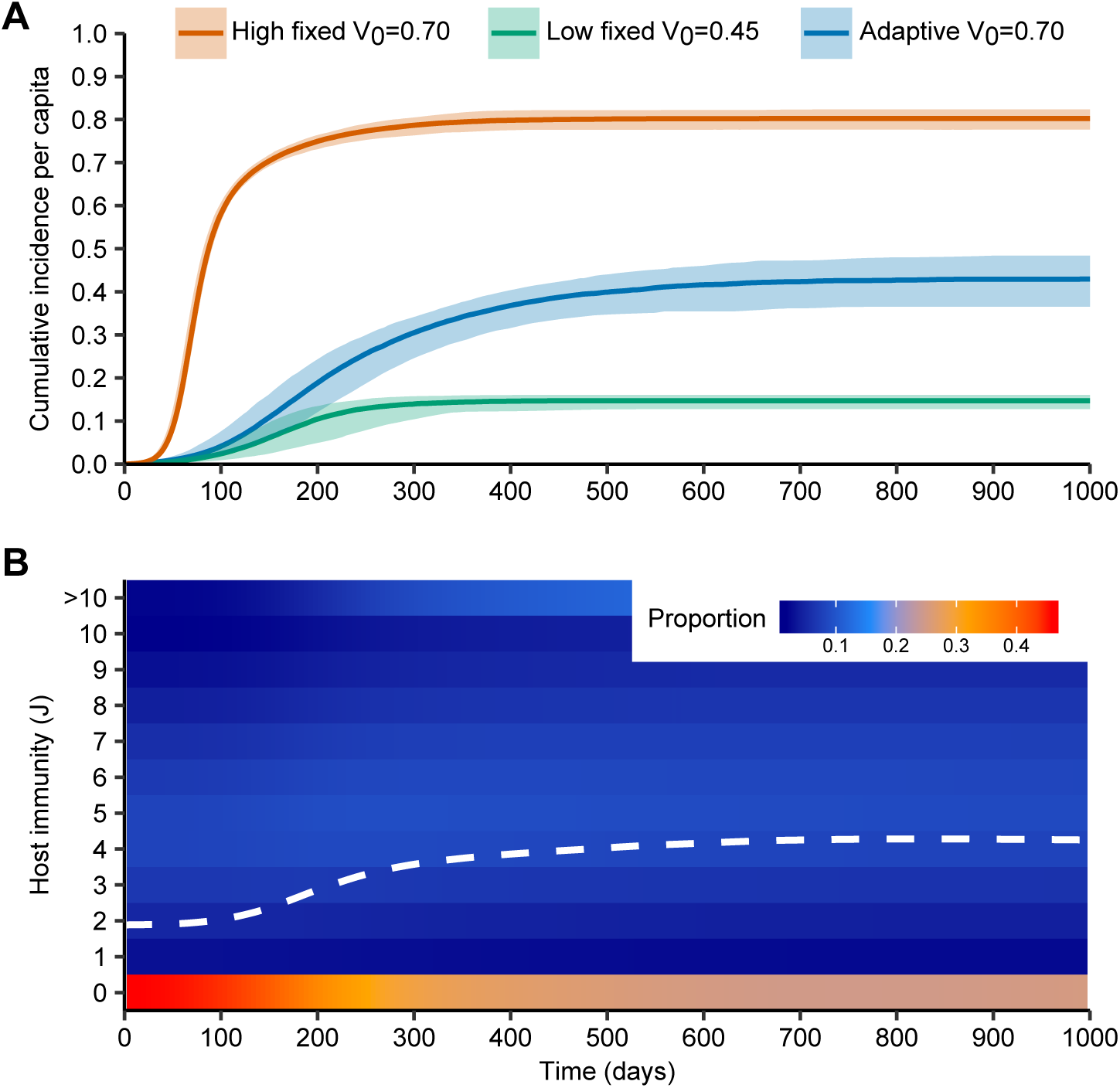
Dynamics of simulated epidemics with and without within-host binding avidity adaptation. (A) Median (lines) and 95% quantiles (shaded regions) of cumulative incidence per capita from 200 simulated epidemics after introduction of an antigenically novel strain into a population size *N* = 500, 000. Blue line shows simulated incidence when binding avidity was allowed to adapt within hosts over the course of the epidemic, starting at a binding avidity of *V*_0_ = 0.70. Orange line shows incidence with binding avidity fixed at values adapted for transmission in a high immunity population *V* = 0.70. Green line shows the incidence with binding avidity fixed at values adapted for transmission in a low immunity population *V* = 0.45. Remaining epidemiological parameters are described in Table S4. (B) Distribution of population partial immunity during the epidemic from one simulation with binding avidity adaptation. Colours show the proportion of susceptible hosts with a given level of host immunity against the seed virus, *J*, over time. White line represents mean population immunity.

To understand the impact of binding avidity adaptation on epidemic dynamics, we compared the simulations with ongoing binding avidity adaptation to simulated epidemics with binding avidity fixed at the lower early-phase adapted value (0.45) and the higher late-phase adapted value (0.70) (Figure 5A). Epidemics with binding avidity adaptation demonstrated higher attack rates relative to simulated epidemics with binding avidity fixed at a lower value (mean of simulations = 43.0%, standard deviation = 2.86% vs. 14.6%, standard deviation = 0.83%). This was due to the gradual increase of binding avidity alongside increasing population immunity, which increased the relative reproductive success of the virus population. However, attack rates under the binding avidity adaptation simulations were lower than in epidemics with binding avidity fixed at the late-phase adapted value (mean of simulations = 80.2%, standard deviation = 1.16%), indicating reduced fitness. This reduction in incidence contradicts the initial hypothesis that binding avidity adaptation should invariably lead to more transmission events during the epidemic.

### Binding avidity adaptation reduces population fitness

We next compared the observed mean binding avidity to the theoretical population-level optimum binding avidity over the course of the epidemic to explain why within-host binding avidity adaptation resulted in reduced transmission relative to binding avidity fixed at the late-phase adapted value. Optimum binding avidity was calculated as the avidity at which the largest number of infected individuals would be produced in the population, as measured by the effective reproductive number *R*_*t*_. During the outbreak, the optimum binding avidity gradually increased to a higher level due to the accumulation of partial immunity in potential hosts. However, we observed that the mean binding avidity of viruses among infected individuals, both before (at the time of infection) and after within-host adaptation (at the end of the infectious period), was lower than the population-level optimum binding avidity and provided lower population-level fitness (Figure 6). In addition, the mean binding avidity measured after within-host adaption (measured at the end of the infectious period) tended to be even lower, corresponding to a lower (*R*_*t*_) than at the start of the infectious period. This could explain why a lower proportion of high net charge isolates and lower average net charge were observed in the external nodes of the previous phylogenetic tree (Table S2).

**Figure 6.**
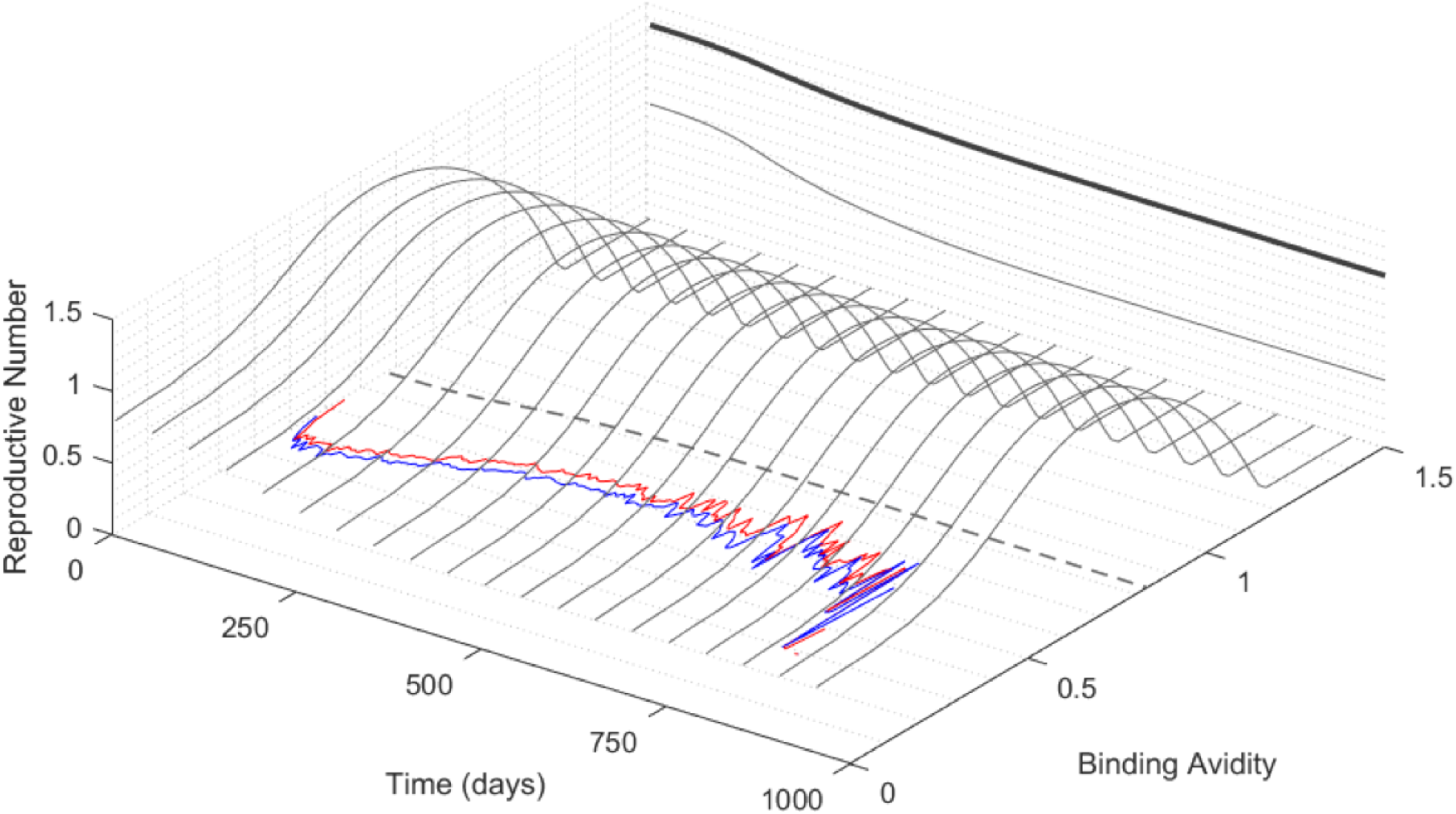
Change of mean binding avidity and effective reproductive number over time. Optimum binding avidity (dotted line) is calculated as the avidity that produces the largest number of infected individuals at the population level, measured by the effective reproductive number *R*_*t*_. The maximum *R*_*t*_ (bold gray line) was produced by the optimum binding avidity whereas the lower *R*_*t*_ (thin gray line) was produced by binding avidity fixed at 0. The red line shows the mean binding avidity across all extant viruses before within-host adaptation (at the start of each individual’s infectious period). The blue line shows the mean adapted binding avidity across all extant viruses after within-host adaptation (at the end of the infectious period). The bell-shaped curves represent *R*_*t*_ at different binding avidities at 50 day intervals.

### Stabilizing selection of binding avidity on the viral phylogeny

Stabilizing selection of binding avidity was demonstrated by comparing the binding avidity values at the internal and external nodes of viral phylogenies reconstructed from our model simulations. While mean binding avidity tended to increase over time, many of the viruses that adapted to much lower binding avidities were present as external tips, which were more likely to die out without onward transmission relative to internal tips (Figure 7 and Figure S4). This suggests a deleterious effect on the virus population. To further understand whether most of the binding avidity changes produced through within-host adaptation were deleterious, we defined a threshold (Δ*V >* 0.3) for significant binding avidity change, and found that external tips in simulated viral phylogenies contained more frequent significant changes (Table S3), similar to the pattern generated by the influenza A/H3N2 sequence data (Table S2). These observations suggest that although many large changes in binding avidity allowed the virus to infect or adapt to more immunologically naive susceptible hosts, the opportunity for the virus to further transmit was lowered in a population with partial immunity. When comparing binding avidity changes between internal (lineages that persisted) and external nodes (lineages that died out) (Figure 8), changes in binding avidity of internal nodes narrowed towards the mean. Nodes in the main lineage were selected to have weaker or near zero binding avidity changes (Figure S7). These results demonstrate a typical pattern of stabilizing selection during an outbreak ([34], [35], [36]).

**Figure 7.**
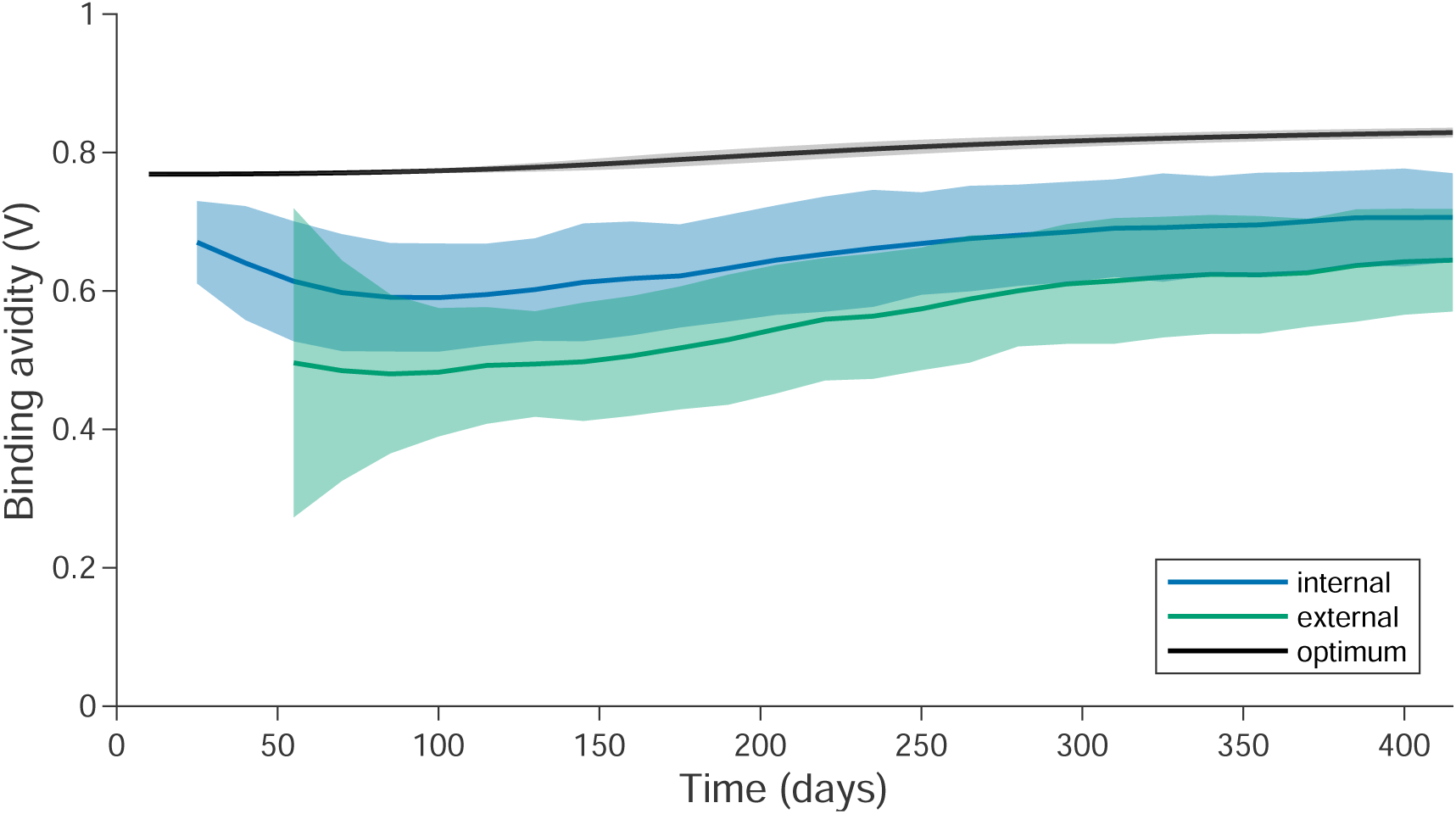
The deleterious effect of within-host binding avidity adaptation on simulated viral phylogenies. The viral phylogenies were produced from the model using 200 simulations. A single viral phylogeny with ancestor nodes was reconstructed from each simulation. Blue represents internal nodes and green represents external tips from the viral phylogeny. Gray represents the optimum virus binding avidity that could produce the highest effective reproductive number *R*_*t*_. Solid line denotes the mean value and the shaded area represents 95% quantiles.

**Figure 8.**
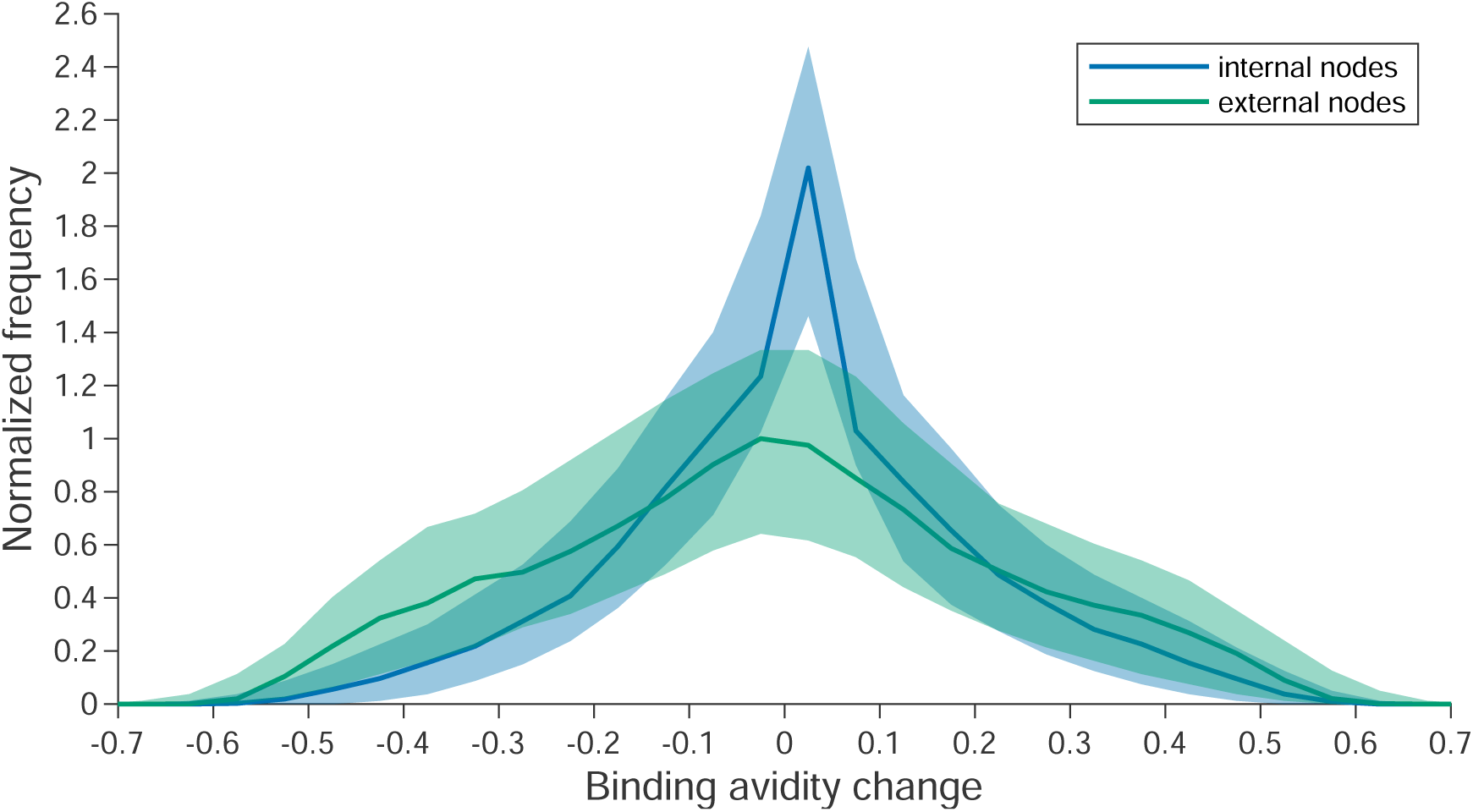
Changes in binding avidity of internal and external nodes (lineages that persisted and died out respectively), averaged from 200 simulated phylogenetic trees. Each simulated tree was randomly thinned to contain 300 external nodes. Y-axis denotes the absolute frequency divided by the mode for external nodes.

## Discussion

Following experimental support for the functional role of cellular receptor binding avidity in immune escape and antigenic drift ([4],[37], [38]), a recent study demonstrated that only a small number of potential substitution sites in the HA head, all near the receptor binding site, are responsible for most of the observed variation in the evolutionary history of influenza A/H3N2 ([22]). Later, deep sequencing studies revealed that potential antigenic variants occur at low frequencies in infected humans ([39]). Together, these findings suggest that although high mutation rates may generate a large number of influenza antigenic variants, many of these variants may not become fixed in a population even when they are expected to have a strong fitness advantage due to immune escape. Furthermore, many recent within-host studies have demonstrated that positive selection from individual-level immunity may play only a limited role in the evolution of seasonal influenza ([20],[39],[40],[41]). These findings raise the question of why within-host immune-mediated selection appears to limit diversification in adaptive evolution of seasonal influenza viruses. We have provided here empirical and theoretical support for stabilizing selection acting on influenza cellular receptor binding avidity in human populations, which may act to limit the number of possible immune escape substitutions. This mechanism may in part modulate seasonal influenza epidemic intensity, despite the fitness advantage that antigenically novel mutants should have at a population level.

Understanding the impact of within-host immune selection pressure on human influenza evolution with a focus on receptor binding avidity requires the characterization of binding avidities using a large number of influenza isolates. Although an RDE assay is useful for determining binding avidity, it is not practically feasible due to prohibitive effort and cost. We found that influenza net charge is correlated with influenza cellular receptor binding avidity and can be used as a proxy for receptor binding avidity based on all of the experimentally derived binding avidity data that we could collect from either the literature or from research groups up to 2017.

Using net charge as a model for cellular binding avidity, the first question we were interested in was whether or not adaptation of influenza binding avidity is present among humans of different ages. Multiple studies have suggested that age-specific immune experience exists, but to our knowledge, no age-specific viral traits have yet been found ([16],[41]). Binding avidity was found to have a strong age effect in our study using net charge as a proxy. Similar patterns between the age distribution of strains with high HA net charge and human seroprevalence have been observed, indicating that adaptation of influenza binding avidity to herd immunity in human population may be present in the evolutionary history of influenza.

Second, we observed that viral net charge evolves slowly with limited diversity in the influenza phylogenetic tree. The selective pressures during an epidemic can be explained as a combination of directional selection acting on a longer time scale (during the epidemic) and stabilizing selection acting on a shorter time scale (during each single infection) using our model simulation results. The selection for a higher mean binding avidity toward a relatively higher effective reproductive number, *R*_*t*_, against the accumulation of population immunity acts as directional selection under a gradually changing environment. However, the diversity of binding avidity will be limited due to a reduced chance for success of a single infection when immune statuses between the infected and contacted persons are not matched. For instance, after an infection in an immunologically naive person, within-host receptor binding can adapt to a very low level in favour of increasing within-host reproductive fitness. However, extreme binding avidity values would often reduce the chance of successful transmission to a randomly contacted person who may have average (relative to the population) partial immunity.

In order to understand the interplay between immune response and receptor binding evolution, we developed an individual-based epidemic model incorporating within-host virus binding avidity adaptation and individual immune boosting. The model allowed us to produce a simulated viral phylogeny and to observe changes of binding avidity alongside changes in individual host immunity over time. Although recent developments of multi-scale, immuno-epidemiological models allow a better understanding of the impact of within-host dynamics on population-level quantities ([42], [43]), partial and stratified immunity, which could influence epidemic dynamics ([11]), have not been fully considered in these models. We have provided here empirical and theoretical support for stabilizing selection acting on binding avidity in human populations with varying degrees of partial immunity.

Our analysis is subject to a number of limitations, and alternative hypotheses exist which challenge some of our assumptions. First, our dataset contained a relatively small number of full-length sequences. Further work should consider alternative, larger datasets to confirm if the pattern of contrasting binding avidity adaptation at external and internal nodes is reproducible. Second, we assumed that within-host immune selection was possible throughout the time course of an infection in our model. Recent theoretical work has proposed a lack of antibody-mediated selection during influenza viral replication because viral growth largely subsides before a humoral antibody response is mounted; antigenic selection instead occurs at the point of transmission via secretory IgA on mucosal surfaces ([44]). The authors suggest that this mechanism can limit population-level diversification because the probability of a novel variant successfully infecting an immunologically experienced host is very low. Our work is not necessarily incompatible with this hypothesis: within-host selection may still act on receptor binding avidity during viral replication whereas selection for immune escape may occur primarily at the point of transmission. The binding avidity of variants that are successful in escaping mucosal immunity may differ to variants that are successful replicators. Third, our simulation results required parameter values to be chosen to reflect within-host dynamics. We chose parameters to qualitatively reflect influenza epidemiology and natural history (e.g. infectious period, distribution of population immunity), but it was not possible to choose all parameters based on empirical data.

Short-sighted evolution of RNA viruses has been observed in recent years, particularly in chronic viral infections such as HIV and HCV, which have long transmission intervals ([45]). Lythgoe *et al.* demonstrated that HIV evolution is “short-sighted” because what is good for the virus in the short-term within the host is not necessarily good for the virus in the long-term at the epidemiological level. Thus, the selective advantage of a viral trait differs at the within-host and between-host level. However, for rapidly evolving viruses which have short transmission intervals, like influenza, it has been suggested that little short-sighted evolution should be possible because there is only a short window of time for within-host adaptation before onward transmission or viral clearance ([46]). Our results suggest that within-host binding avidity adaptation to a very low value or a very high value in individuals with immunologically naive or strong partial immunity becomes detrimental to the survival of a viral lineage in a population with pre-existing partial immunity. Binding avidity adaptation as a trait may therefore be selected against to reduce short-sighted evolution. Whether the short infection duration of influenza coupled with a high transmission rate is a consequence of short-sighted evolution is an open question.

Deleterious mutations have been proposed to affect influenza evolution and antigenic drift ([47], [48]). This mechanism would require a sufficiently large number of mutations with detrimental effects. Mutations that affect protein stability can be produced ([49]), but given a high mutation rate and rapid rate of evolution within a host, the percentage of deleterious mutations would be expected to be low due to positive selection. Our results are not inconsistent with the proposed effects from deleterious mutations. Because partial and stratified immunity develops during an epidemic, many immune escape mutations led by binding avidity changes within a host will reduce the effective reproductive number at the population level and behave like deleterious mutations due to the trade-off between adaptation at the within and between host scales. This explains why many antigenic or immune escape substitutions may not become fixed in a population even when they would be expected to have a strong fitness advantage [39].

Optimal vaccination strategies can be better assessed by considering the impact of receptor binding adaptation on transmission in a heterogeneous population. This study highlights the potential benefit of incorporating binding avidity assays in addition to serological tests in order to prioritize the groups that should be vaccinated to reduce overall disease incidence and understand constraints on influenza antigenic evolution or antigenic drift ([50], [14]). Understanding the effect of binding avidity on both epidemic and evolutionary dynamics would be beneficial for influenza outbreak future prediction and control.

## Material & Methods

### Analysis of HA cellular receptor binding avidity

The RDE binding avidity for amino acid changes were collected from the literature ([4] [6] [5] [38]) and also obtained directly from a group performing RDE assays ([4]). In total, 74 mutant strains, including 71 from A/H1N1 and 3 from A/H3N2, were collected (Table S5). After removing 3 duplicates and 1 reversed mutation, there were 70 mutants, including 49 single, 20 double and 1 triple amino acid changes. For the RDE binding avidity assays present in the literature, the magnitude of each change was extracted using the Web Plot Digitizer ([51]). The effect of each single amino acid change (including secondary and tertiary changes) on binding avidity was calculated as the log ratio of RDE activity of the mutant compared to the wild type. Binding avidity against the actual net charge of HA from each mutant strain was calculated as the log ratio of RDE activity of the A/H1N1 mutants (n=67) relative to wild type influenza A virus (strain A/Puerto Rico/8/1934 A/H1N1), also called PR8.

### Calculating net charge of influenza HA

Influenza virus sequences isolated from New York State as part of the Influenza Genome Sequencing Project ([27]) along with the exact date of isolation and clinical metadata on host age were collected from the Influenza Virus Resource Database ([28]). Among total 759 isolates, we used 686 full-length influenza A/H3N2 HA sequences, isolated between July 1993 and June 2006. Changes in net charge relative to reconstructed ancestral sequences were estimated by summing the total number of positively charged amino acids minus the number of negatively charged amino acids. Amino acid such as Arginine (R), Histidine (H) and Lysine (K) are positively charged whereas Aspartic acid (E) and Glutamic acid (D) are negatively charged. For estimating net charge from the isolated viral HA sequences, net charge was calculated from the sequences using amino acid positions 17 through 345 of the HA, which define the protein’s globular head domain (HA1). We specifically did not calculate net charge values using only the amino acid positions that comprise the cellular receptor binding site because charge at other sites in the globular head domain still impact electrostatic forces and therefore binding avidity.

### Phylogenetic Analysis

Phylogenetic trees were reconstructed using the software BEAST ([52]), under a general time reversible (GTR) model with gamma distributed rate variation and a proportion of invariant sites. 10^7^ Markov chain Monte Carlo (MCMC) steps were sufficient in reaching high effective sample sizes (*>*100). From each viral subtype’s maximum clade credibility (MCC) tree, we inferred the ancestral states of the observed viral sequences. If BEAST produced multiple possible ancestor sequences, consensus sequences were calculated with a BLOSUM50 scoring matrix using Matlab Bioinformatics Toolbox. For all the sequences at internal and external nodes we calculated net charge as described above.

### Individual-based model

We developed a stochastic Susceptible-Infected-Recovered-Susceptible (SIRS) individual-based model embedded with within-host receptor binding adaptation and immune responses to track individual human hosts and viruses ([11]). The status of each individual host could be changed from *S* to *I, I* to *R* with short-term full protection with respect to infection, and from *R* to *S* with partial immunity *J* (Figure S5A). At the time of contact between a susceptible and infected host, the probability of infection *ρ* was determined by host immunity against the challenging source virus and its binding avidity as described below. Properties of viruses, including receptor binding avidity changes *V* and antigenic distance *δ*, were tracked within hosts and between each transmission event (Figure S5B). For computational reasons, we simulated one virus object per infection, such that each virus represented the average of all individual virions in that virus population during infection in a host. All epidemiological parameter definitions and assumed values, including births and deaths at rate *µ*, the infectious period 1*/γ*, short-term full immunity that wanes at rate *ω*, contact rate *c*, etc., are shown in Table S4.

The model and code are openly available and documented as an R-package with a C++ back-end at https://github.com/jameshay218/driftSim. The R-package allows the exploration of a number of epidemiological mechanisms that we do not consider in these analyses (e.g., antigenic drift, antigenic change as a by-product of binding avidity adaptation, multi-season dynamics). We restricted the analyses here to simplified use cases of the R-package as described below. The details of how simulation was carried out are described in the Supplementary Methods.

### Stratified host immunity

In the model, host immunity determined the probability of success of two events: (i) infection during a transmission event and (ii) within-host replication during the course of an infection. Host immunity was determined by the contents of an individual’s infection infection history, **h**, and their homologous antibody titer, *j*. We defined an infection history **h** as the set of all *n* viruses that an individual had previously been infected with:

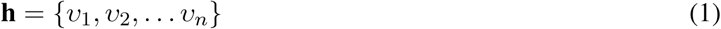

where *v*_*n*_ represents virus *n*. When calculating the probability of infection during a contact (the first event), the effective immunity of a susceptible host *i* against infection from the source virus in an infectious host *s* was given by the total antibody titer less the smallest antigenic distance of the source virus *v*_*s*_ (i.e., from host *s*) to all viruses in the susceptible host’s infection history **h**_*i*_:

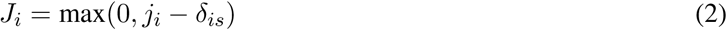

where *δ*_*is*_ = min(*δ*_*ls*_) for *l* ∈ **h**_*i*_; and *j*_*i*_ is the homologous titer of host *i* against any strains that they had previously been infected with (Figure S5A).

Determining host immunity to calculate the within-host reproductive success of a virus (the second event) was defined in a similar way: the measure of host immunity for an infected host *i* was *J*_*i*_ = max(0, *j*_*i*_ − *δ*_*i*_), where *δ*_*i*_ is the smallest antigenic distance of the virus infecting host *i* relative to all viruses that previously infected host *i* (i.e., antigenic distance to self) (Figure S6).

In these analyses, we assumed that only one initial antigenically novel virus strain was seeded in 100 initially infected individuals (*I*_*t*=0_ = 100) and no further antigenic mutations occurred during the epidemic. All individuals in the initial population were assumed to have the same infection history such that *δ*_*is*_ = *δ*_*i*_ = 3 for all viruses and hosts at the start of the simulation. Thus, all instances of *δ*_*is*_ and *δ*_*i*_ were either 0 or 3 depending on whether the host had been previously infected with the novel virus or not. Similarly, infection histories could be either **h** = ∅ (empty infection history at birth or the start of the simulation) or **h** = *{v*_1_*}* (infected with the seed virus one or more times). However, individuals could have different pre-existing immunity. *J* can be considered analogous to immunity as determined by antibody titers ([32] [11]). The definition of antigenic distance used here is analogous to previous definitions ([1]).

### Antibody boosting

We assumed that recovery generated an antibody boost drawn from a Poisson distribution ([11]) and that the level of realised boosting was inversely proportional to the pre-existing antibody titer, representing a ceiling effect on antibody responses ([53]). Antibody boosting was therefore given as:

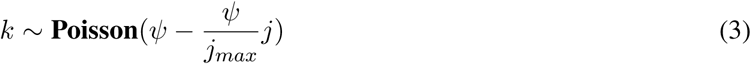

where *j* is the antibody titer before boosting; *ψ* is the mean antibody boost from a starting titer of 0; and *j*_*max*_ is the maximum achievable antibody titer. The boosted antibody titer following recovery was therefore given as *j*′ = *j* + *k*. Figure S6 shows an example of how an individual host’s immunity changes over time following single or multiple infections. Hosts could be infected any number of times, but the probability of re-infection decreased as host immunity increased following recovery.

Similarly, once host *i* recovered from infection, that virus was added to *i*’s infection history such that:

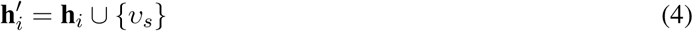

### Probability of infection

In the stochastic model, each time a source virus came into contact with a susceptible host, the probability of successfully infecting that host was defined as:

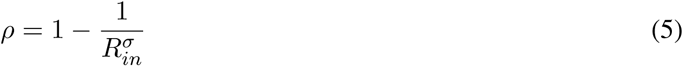

where *R*_*in*_ was the within-host reproductive number and *σ* was the effective number of infectious particles initially transmitted. Note that although the effective number of infectious particles can be *>* 1, this number only affects the probability of a single infection event during transmission and only one virus object was simulated for each infection. Following our previous study ([14]), *R*_*in*_ was defined as the expected number of virions produced by a virus in a host with immunity *J* :

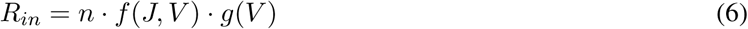

where *n* is the effective number of virions per replication; *f* (*J, V*) = [1 − *e*^*p*(*V* +1)^]^*rJ*^ gives the probability of an infecting virus with binding avidity *V* evading the immune response given host immunity *J* as defined above; 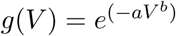 gives the probability of successful replication within a host with binding avidity *V* ; and *p, r, a* and *b* are scaling constants ([14]). Combining these probabilities with the effective number of virions per replication gave the within-host reproductive number of viruses.

### Within-host receptor binding adaptation

During the infectious period, binding avidity *V* of each infectious virus adapted to the host’s immunity *J*. The change of binding avidity with time Δ*V* (*t*) was defined as the product of the rate of change of the virus binding avidity given by *dV/dt* = *k*_*c*_(*dR*_*in*_*/dV*) and time Δ*t* ([14]), *where dR*_*in*_*/dV* was the fitness gradient with respect to binding avidity, and *k*_*c*_ quantifies the amount of genetic variance in receptor binding avidity within a single host as a constant mediating adaptation rate.

### Initial population immunity

We began each simulation using initial conditions for the immune profile of susceptible hosts produced after simulating a single epidemic peak to represent a population with realistic heterogeneity in pre-existing immunity. To match the distribution of partial immunity that is observed in human populations, we ran the simulation from a completely naive population to generate a distribution of host immunity. This initial-condition generating epidemic generated an attack rate of 54%. Each infected individual received a boost to immunity drawn from a Poisson distribution with mean *ψ* = 8 following infection, resulting in a distribution of host immune states, *J*. All simulations were then run using the same host immunity distribution and introducing a new virus with antigenic distance *δ* = 3 relative to the strain used for the generation of initial conditions. The resulting overall distribution of effective immunity *J* was similar to observed serological data for seasonal influenza ([11]).

### Computational analysis

Analyses were performed using a combination of R version 3.5.0 (R Foundation for Statistical Computing, Vienna, Austria) and Matlab. R scripts and Matlab code to reproduce all analyses are available at https://github.com/hy39/bindingAvid and https://github.com/jameshay218/driftSim.

## Acknowledgements

We thank Jonathan Yewdell for giving valuable suggestions and Scott Hensley for providing RDE binding avidity data. We also thank Katia Koelle and Christophe Fraser for valuable suggestions. This study was supported by grants from the City University of Hong Kong (#7200573 and #9610416).

## Supplementary

**Supplementary Methods. Further description of individual-based model implementation.**

**Table S1.**
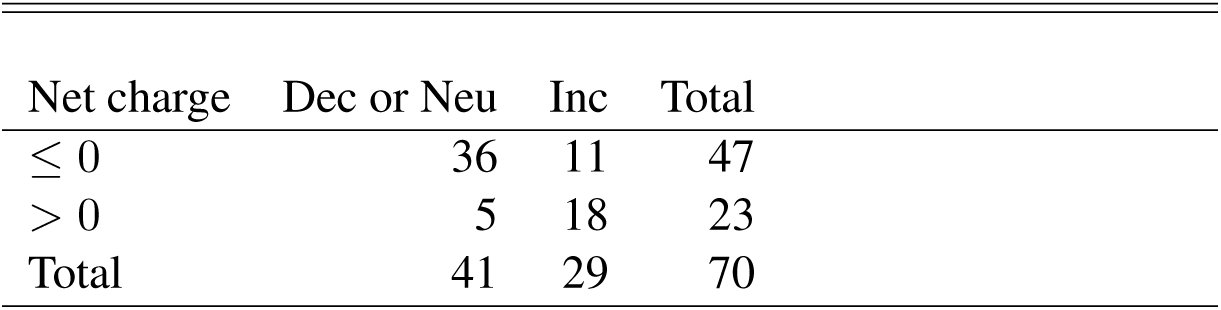
Change in binding avidity from HA mutations by net charge category. The odds ratio of increasing receptor binding avidity for mutations with positive compared with non-positive net charge change was given. (Odds ratio = 11.27; 95% confidence interval: 3.14-48.5; p-value *<*0.001 using Fisher’s exact test). *Dec or Neu* indicates decreased or neutral effects on receptor binding avidity and *Inc* indicates increased effects on receptor binding. 70 mutations were used after removing 3 duplicates and 1 reversed mutation.

**Table S2.** Comparison of the net charge distribution of internal and external nodes on the phylogenetic tree. The difference of the number of strains that changed net charge (*# Without net charge change* and *# With net charge change*) between external nodes and internal nodes was tested using Fisher’s exact test, giving an odds ratio (odds of higher number of net charge changes at external nodes relative to internal nodes) of 1.64 (95% CI: 1.14-2.35; Fisher’s exact test; p-value=0.009; n=1370). The number of internal (or external) nodes that fit this definition is expressed as a percentage of the total number of internal (or external) nodes. Among the nodes with net charge change, the number of positive *Pos* and negative *Neg* changes are shown. High net charge is defined as an absolute net charge greater than 18 (median of all 1370 samples is 17). The average net charge and standard deviation of internal and external nodes is shown.

|  | # Without<br>net charge<br>change | # With net<br>charge<br>change | # Pos<br>changes | # Neg<br>changes | Prop high<br>net charge<br>(%) | Avg net<br>charge |
| --- | --- | --- | --- | --- | --- | --- |
| Internal | 631 (92.3%) | 53 (7.7%) | 15 | 38 | 20.0 | 17.52 ± 1.06 |
| External | 603 (87.9%) | 83 (12.1%) | 32 | 51 | 18.7 | 17.45 ± 1.09 |

**Table S3.** Comparison of the number of nodes with and without a significant change in binding avidity. Binding avidity changes were calculated by comparing each node’s binding avidity relative to their direct ancestors’. A node’s binding avidity was defined to have a significant change when the difference from its direct ancestor’s binding avidity exceeded a threshold of 0.3 in Δ*V*, resulting in approximately 20% of nodes demonstrating a significant change in binding avidity. Results shown are averaged over 200 simulated phylogenetic trees. Each simulated tree was thinned to give 300 viruses as external nodes. Taking equal weights, the harmonic mean p-value ([54]) calculated from individual Fisher’s exact test on each simulation was *<*0.001. Odds ratio from Fisher’s exact test on the simulation means was 2.36 (95% CI: 1.50-3.74; p-value *<*0.001).

|  | Without significant change in<br>binding avidity | With significant change in<br>binding avidity |
| --- | --- | --- |
| Internal | $261.5 \pm 6.5$ ($87.7 \pm 2.2\%$) | $36.5 \pm 6.5$ ($12.3 \pm 2.2\%$) |
| External | $225.4 \pm 9.1$ ($75.1 \pm 3.0\%$) | $74.6 \pm 9.1$ ($24.9 \pm 3.0\%$) |

**Table S4.** List of parameters used in individual-based model simulations.

| Parameter name | symbol | values |
| --- | --- | --- |
| Total human population size | $N$ | 500,000 |
| Birth & death rate (per day) | $\mu$ | $1/(70*365)$ |
| Recovery rate (per day) | $\gamma$ | 1/3.3 |
| Waning rate (per day) | $\omega$ | 1/25 |
| Initial number of seeds | $I_{t=0}$ | 100 |
| Antibody boosting | $\psi$ | 8 |
| Host immunity | $J$ | varied |
| Initial antigenic mutant distance | $\delta$ | 3 |
| Effective number of transmitted virions | $\sigma$ | 1 |
| Effective number of virions per replication | $n$ | 4 |
| Contact rate (per day) | $c$ | 0.7 |
| Binding avidity adaptation rate (per day) | $k_c$ | 0.3 |
| Maximum achievable host immunity | $j_{max}$ | 40 |

**Table S5.** List of all collected mutations with corresponding binding avidity (RDE). (Attached as an excel file)

**Figure S1.**
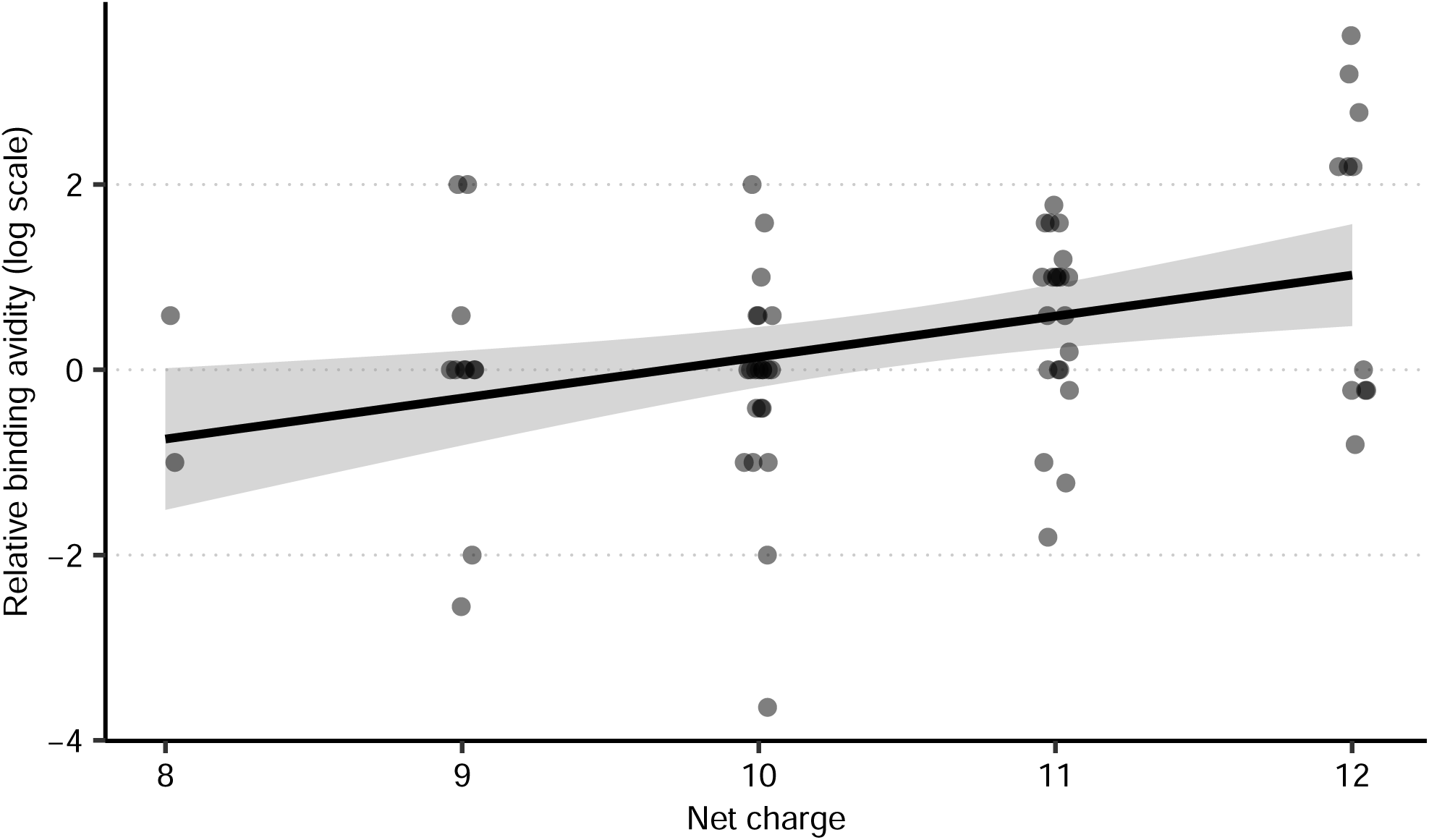
Relative binding avidity against net charge of A/H1N1 HA. Relative binding avidity was calculated as the log ratio of RDE activity of the mutant relative to the wild type A/H1N1 PR8. Net charges of all 67 A/H1N1 mutant strains were calculated based on their amino acid sequences using positions 18 to 342 of HA1. Pearson correlation coefficient was r=0.353 (95% confidence interval: 0.123-0.547; p-value=0.003) and slope of the line was 0.282 (p-value=0.002 using a one-tailed test to test whether the slope was greater than zero).

**Figure S2.**
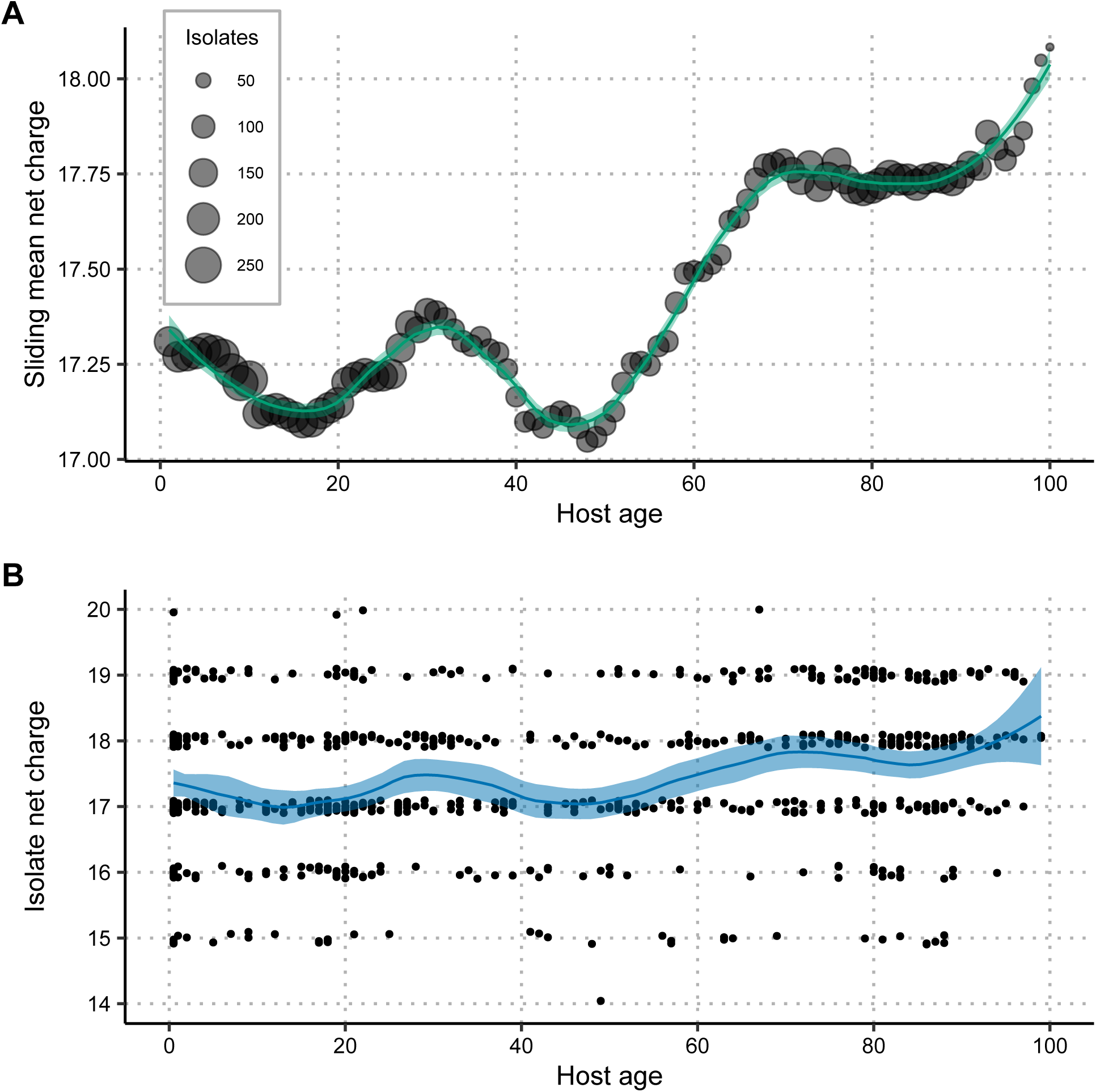
Virus net charge distribution by age of host. (A) Sliding mean net charge by age. Each circle represents the mean net charge calculated from a sliding window of size 10 centered on each year from 1 to 100 yo. The size of circle represents the number of strains in a given sliding window. Green line and shaded region show a LOESS smoothing curve fit with 95% confidence interval (CI) to the calculated values using a span value of 0.3. (B) Isolate net charge by age of host. Each point is one virus isolate slightly jittered in the y-axis. Blue line and shaded region show LOESS smoothing curve fit with 95% CI using a span value of 0.3. 759 virus sequences and age metadata were obtained from the Influenza Genome Sequencing Project ([27]) and the Influenza Virus Resource Database ([28]).

**Figure S3.**
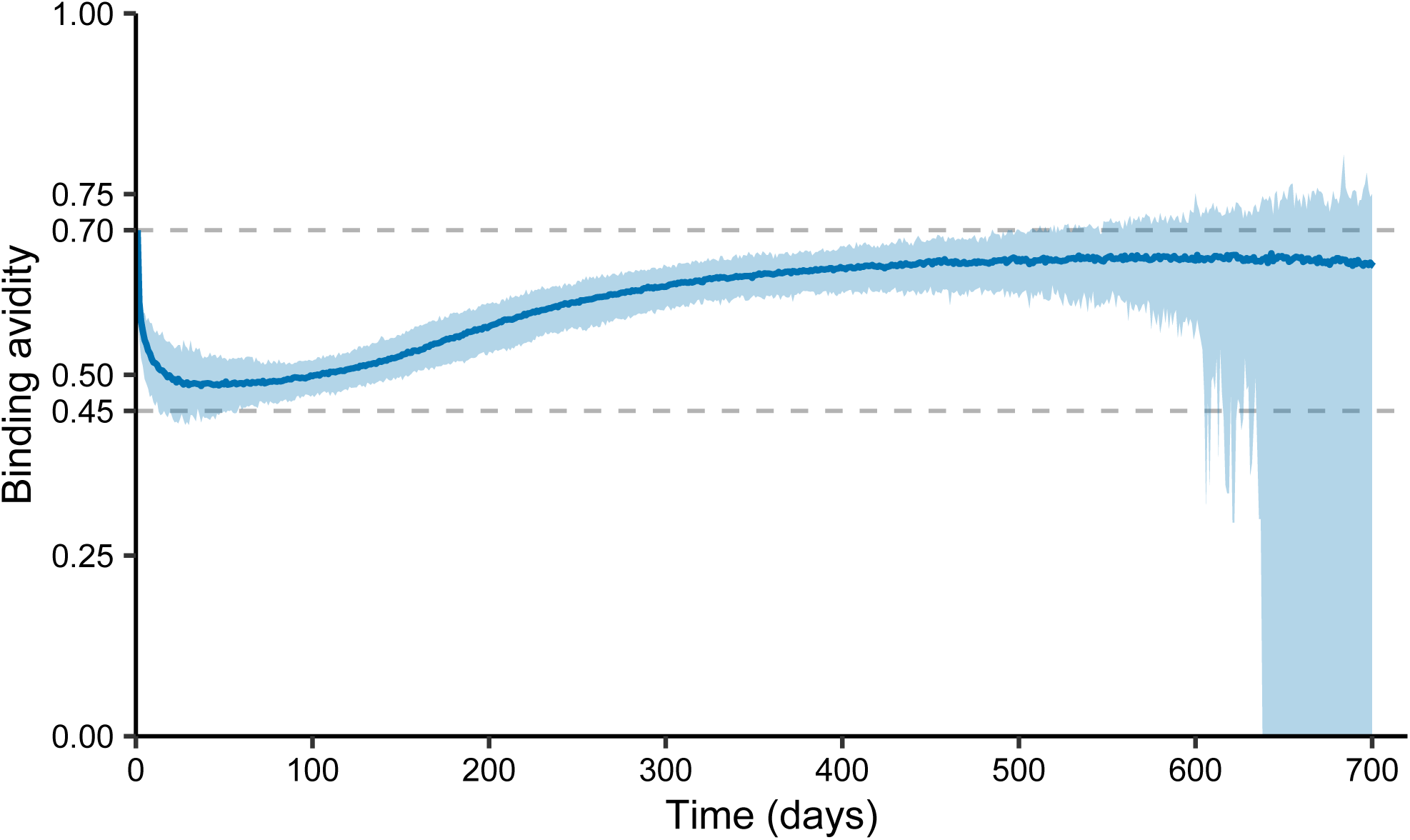
Change in mean binding avidity of all extant viruses over the course of simulated epidemics. Dark blue line and light blue region show median and 95% quantiles for the estimated means produced by 200 model runs in Figure 5. Binding avidity was recorded as 0 once there were no more infected individuals.

**Figure S4.**
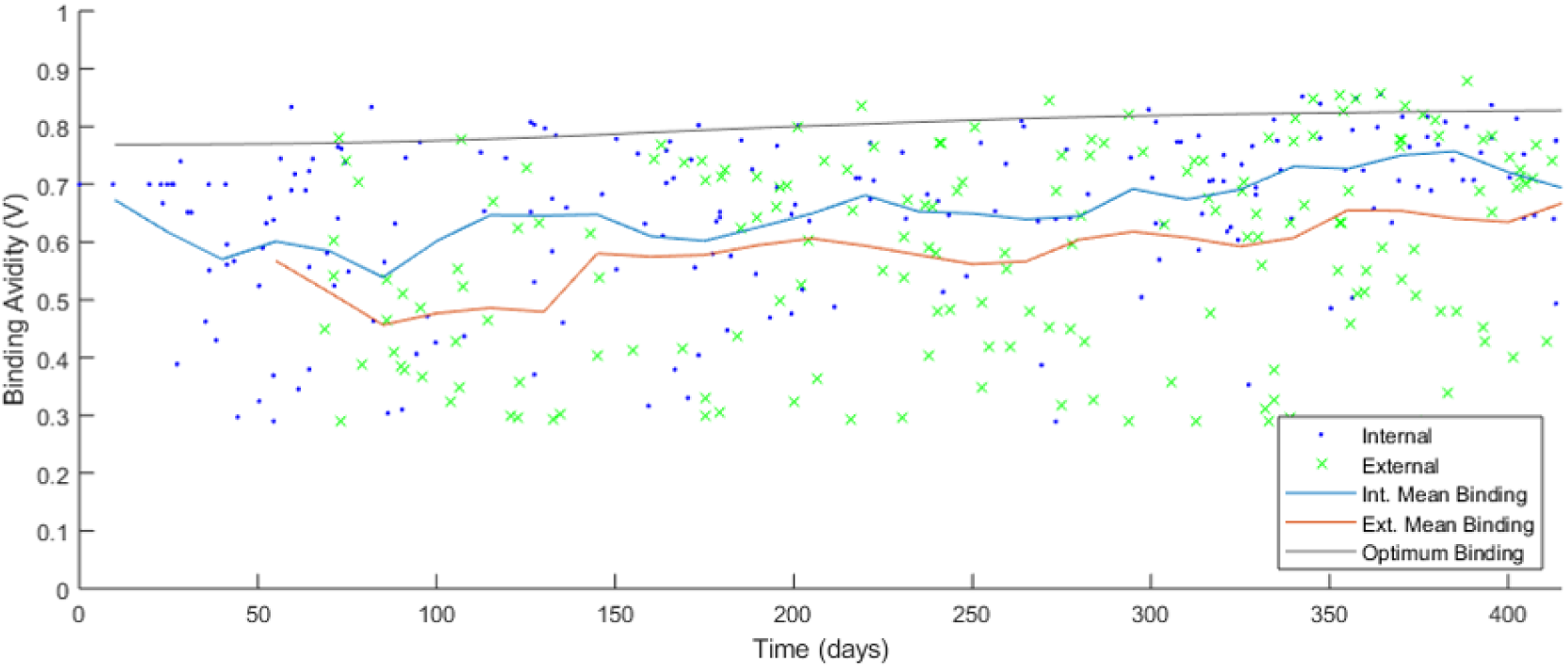
Binding avidity over time in a typical model simulation. Parameter values are the same as those for Figure 4. Blue dots represent internal nodes from the viral phylogeny. Green dots represent external tips from the viral phylogeny. Blue line represents mean binding avidity from internal nodes. Red line represents mean binding avidity from external nodes. Gray line shows the optimum virus binding avidity that will produce the greatest number of infections at the population level.

**Figure S5.**
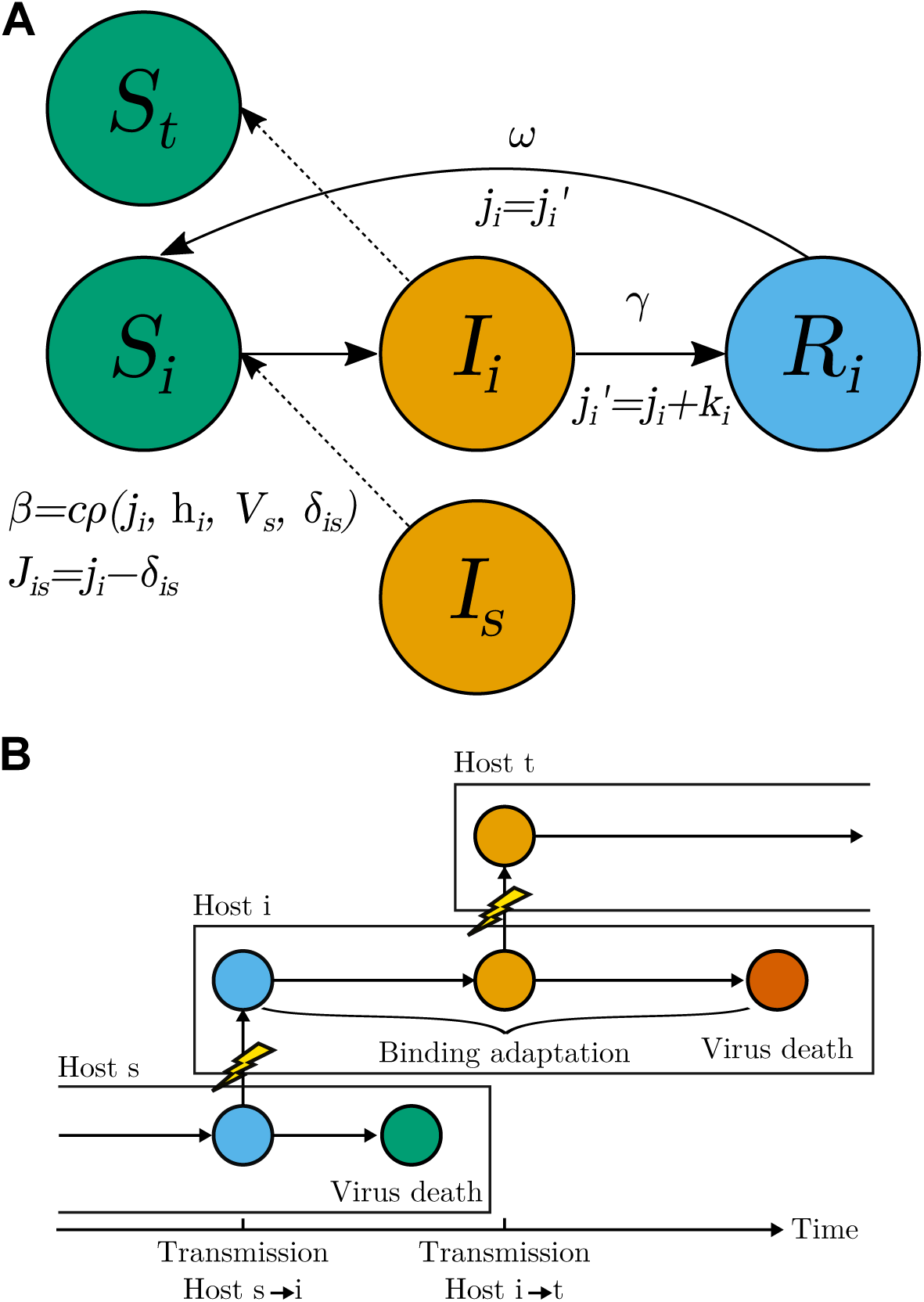
Schema of disease transmission in the individual-based model. The schema represents infections from host *s* to *i* then to *t*. (A) The status of a host during an infection is shown as an SIRS model (S=Susceptible; I=Infected; R=Recovered). *j*_*i*_ is the immunity of the host *i* against all previous infecting viruses. *J*_*is*_ is the immunity of the susceptible host *i* against infection from the infected source *s. δ*_*is*_ is the smallest antigenic distance from the virus affecting host *i* to viruses in the infection history of host *i*, **h**_*i*_. *V*_*s*_ is the binding avidity of the virus from the source *s* at time of transmission. *k*_*i*_ is the antibody boost upon recovery and *j*′ represents the boosted immunity. Epidemiological parameters: *c* is the contact rate, *ρ* is the probability of infection (shown here as a function of host and virus properties), *γ* is the recovery rate, and *ω* is the waning rate of fully protective transient immunity. (B) Viruses with different receptor binding avidities are tracked within and between all individual hosts. Each rectangle represents the infection period of a particular host. After transmission (lightening symbols), a virus is produced with the same binding avidity as it had in the previous host at the time of transmission (blue and orange circles). Over the course of the infection, binding avidity adapts to the host’s immunity level. Circle colors represent changes in virus binding avidity over the infectious period. The virus dies upon host recovery.

**Figure S6.**
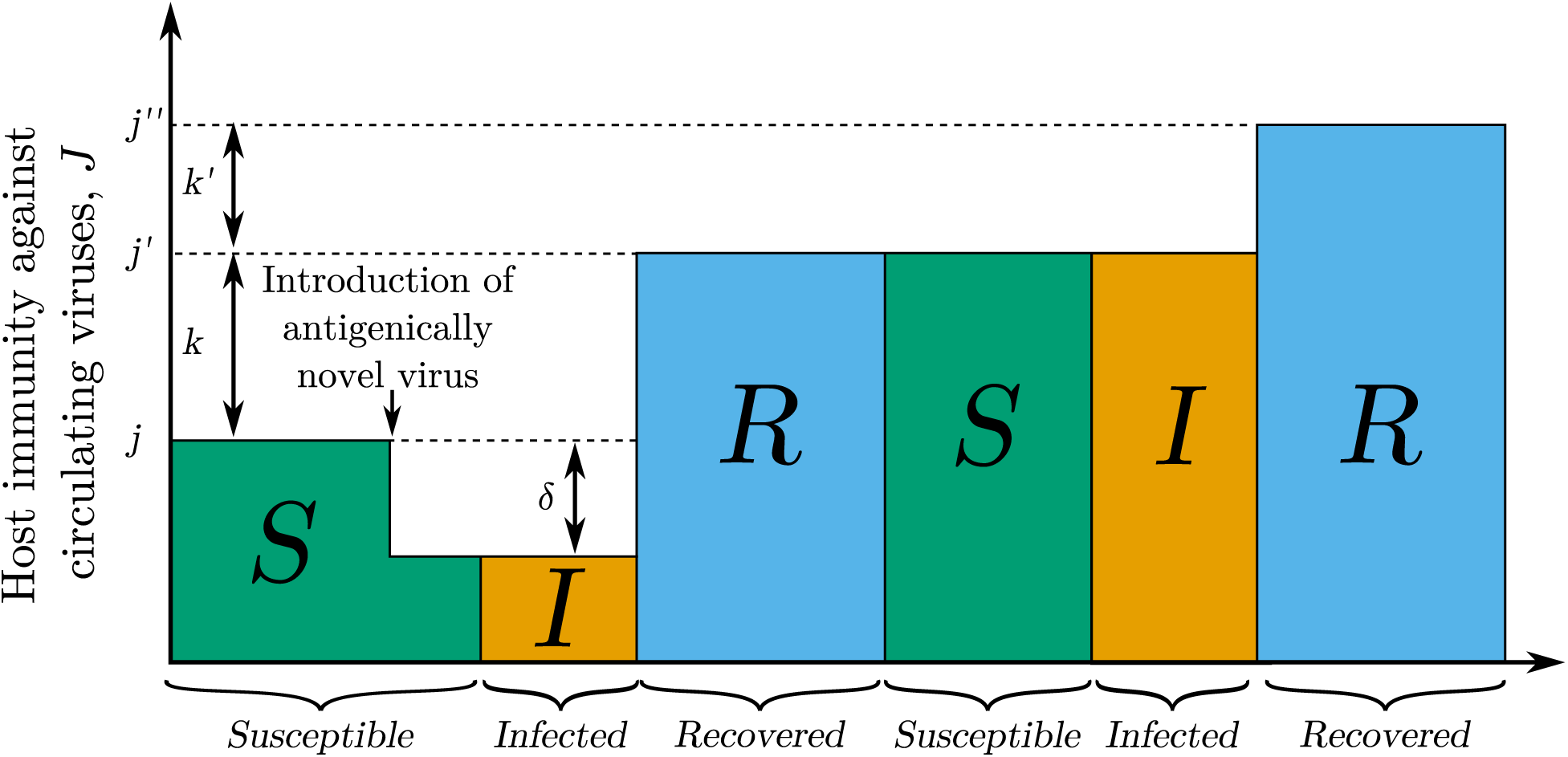
Schema of an individual host’s immunity over time. Colored rectangles show host infection status over time (note that the width of each state does not represent actual time spent in that state). *j* represents the antibody titer available against a virus with no antigenic distance to any virus that the host has been infected with previously (*δ* = 0). If an antigenic mutant emerges such that the shortest antigenic distance between that virus mutant and the viruses that previously infected the host is *δ*, then the host has reduced immunity (*J* = *j* − *δ*) upon contact. The marked point shows the drop in immunity *J* against circulating viruses upon introduction of an antigenically novel virus into the population. Once the host recovers from infection, host immunity to previously encountered viruses *J* is boosted to a higher level from antibody boosting *k* (*j*′ = *j* + *k*) and the virus is added to the host’s infection history **h**. The host then has full immunity (*J* = *j*′) against all viruses in the infection history, including the recent mutant (*δ*=0). If the host is reinfected with the same virus, host immunity is boosted again (*j*″ = *j*′ + *k*′).

**Figure S7.**
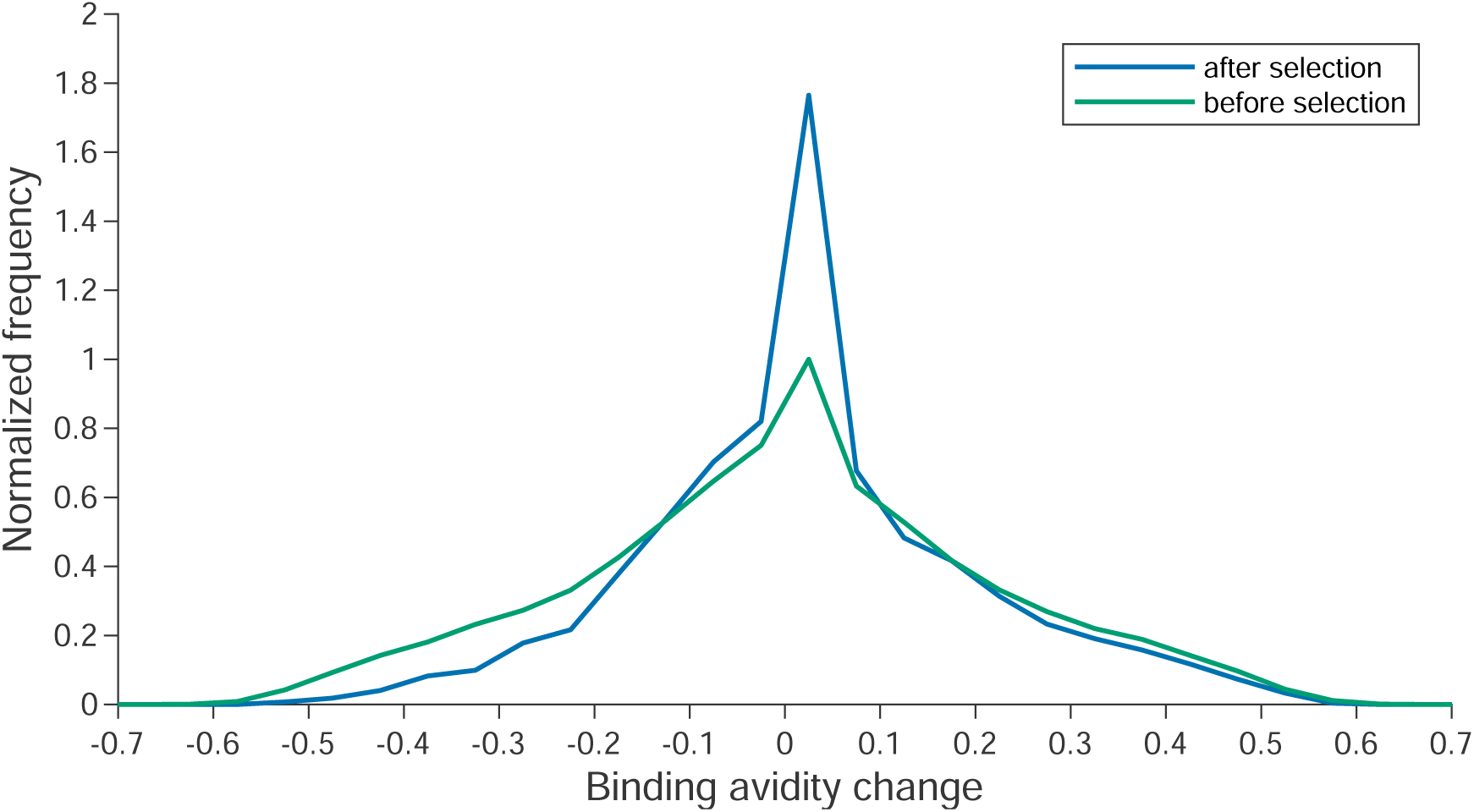
Changes in binding avidity of nodes before (defined as all nodes) and after (defined as internal nodes in the main trunk only) selection, averaged from 200 simulated phylogenetic trees. Y-axis denotes the absolute frequency divided by the mode for nodes before selection.

## Notes

https://github.com/hy39/bindingAvid

https://github.com/jameshay218/driftSim

### Competing Interest Statement

The authors have declared no competing interest.

