## Supplementary Methods 1 for "Stabilizing selection of seasonal influenza receptor binding in populations with partial immunity"

Supplementary Methods: details of individual-based model simulations

### Individual-based model implementation

The model was simulated under demographic stochasticity using Gillespie's tau-leap algorithm ([1]) to calculate the probability of events occurring for each individual in each day. We assumed that births of new hosts occurred at a rate 1/70 per year, and that each new host began with completely naive immunity ($j = 0$ and $\mathbf{h} = \emptyset$). Similarly, deaths occurred at the same rate (1/70) for the entire population thereby ensuring a constant population size. Total number of contacts was calculated based on the contact rate $c$, number of susceptibles $S$, and number of infected individuals $I$ in a population of size $N = 500,000$, resulting in $\frac{cSI}{N}$ total contacts per unit time. For each contact, a random susceptible target $S$ and an infected source $I$ were drawn from the population. The probability of a successful transmission was a function of the properties of the susceptible host's immunity and the infecting virus as described in the main text.

Once an individual became infected, the average infectious period $1/\gamma$ lasted 3.3 days, during which the virus' binding avidity $V$ adapted to the host immunity $J$ at a rate determined by the fitness gradient ([2]). After the infected individual recovered, the individual became transiently fully protected for an average of 25 days ($1/\omega$) through short-lived immunity, possibly mediated by cytotoxic T lymphocytes along with other immunological factors ([3] [4]).

The SIR model is described by the following set of equations:

$$\begin{aligned}
\frac{dS}{dt} &= \mu N - \beta SI - \mu S + \omega R && (1)\\
\frac{dI}{dt} &= \beta SI - (\mu + \gamma)I && (2)\\
\frac{dR}{dt} &= \gamma I - (\mu + \omega)R && (3)
\end{aligned}$$

where $S$ is the number of susceptible individuals; $I$ is the number of infected individuals; $R$ is the number of fully immune individuals; $\mu$ is the daily birth and death rate; $\beta = c\rho$ is the transmission rate as described in the main text; $\omega$ is the rate at which full, short term immunity wanes; and $\gamma$ is the recovery rate.

The model can be approximately described by a set of difference equations which were implemented via the "$\tau$-leap" algorithm to provide the stochastic, agent based form of the above SIR model:

$$\begin{aligned}
S(t + \delta t) &= S(t) + \delta M_B + \delta M_W - \delta M_T - \delta M_{D_S} && (4)\\
I(t + \delta t) &= I(t) + \delta M_T - \delta M_{D_I} - \delta M_R && (5)\\
R(t + \delta t) &= R(t) + \delta M_R - \delta M_W - \delta M_{D_R} && (6)
\end{aligned}$$

where the nomenclature of $\delta M_X$ describes the number of events of type $X$ that occurred within a small, fixed time interval, $\delta t$. The number of events that occurred were approximately Poisson distributed such that:

$$\begin{aligned}
\delta M_B &\approx \text{Poisson}(\mu N \delta t) && (7)\\
\delta M_T &\approx \text{Poisson}(\beta SI \delta t) && (8)\\
\delta M_{D_S} &\approx \text{Poisson}(\mu S \delta t) && (9)\\
\delta M_{D_I} &\approx \text{Poisson}(\mu I \delta t) && (10)\\
\delta M_{D_R} &\approx \text{Poisson}(\mu R \delta t) && (11)\\
\delta M_R &\approx \text{Poisson}(\gamma I \delta t) && (12)\\
\delta M_W &\approx \text{Poisson}(\omega R \delta t) && (13)
\end{aligned}$$

### Binding avidity adaptation

The binding avidity of a given virus, $V_i$, was assumed to adapt to the level of immune selection pressure elicited by the host during the course of infection. We assumed that in a given time step, $\delta t$, during infection, binding avidity changed proportional to the fitness gradient at the current binding avidity level. We defined the fitness gradient to be proportional to the derivative of the within-host reproductive number, $R_{in}$, with respect to binding avidity. We assumed that the binding avidity trait moved "towards" the value that gave the greatest $R_{in}$ over time. Based on $f(J, V_i)$ and $g(V_i)$ as defined in the main text, the rate of change of binding avidity with respect to time is given by:

$$\frac{dV}{dt} = \frac{dR_{in}}{dV} = g(V_i)f'(J, V_i) + g'(V_i)f(J, V_i) \tag{14}$$

where $J = j - \delta$ is defined in the main text.

To avoid solving this equation numerically for every time step for each adapting virus, we pre-computed a table of expected binding avidity change for a set of given binding avidities and host immunity values within a small time step, $\delta t$. After each time step the new binding avidity of a given virus was calculated as:

$$V_{new} = V_{cur} + k_c o(\delta t, V_{cur}, J) \tag{15}$$

where $o(\delta_t, V_{cur}, J)$ was found by solving the ODEs above for the time step $\delta t$ given a current binding avidity value $V_{cur}$ and level of effective host immunity $J$ as described above. $k_c$ was a constant of proportionality.

We approximated these continuous traits by pre-computing the amount of change in time step $\delta t$, $o$, for a set of discrete values of $J$ and $V$. These values were stored in a matrix $O$:

$$O = \begin{bmatrix} o_{V_1,J_1} & o_{V_1,J_2} & o_{V_1,J_3} & \cdots & o_{V_1,J_n} \\ o_{V_2,J_1} & o_{V_2,J_2} & o_{V_2,J_3} & \cdots & o_{V_,J_2n} \\ \vdots & \vdots & \vdots & \ddots & \vdots \\ o_{V_d,J_1} & o_{V_d,J_2} & o_{V_d,J_3} & \cdots & o_{V_d,J_n} \end{bmatrix} \tag{16}$$

where:

$$V_{d+1} = V_d + \delta V \quad (17)$$

$$J_{n+1} = J_n + \delta J \quad (18)$$

$$J = J_n \text{ if } J_n \leq J < J_{n+1} \quad (19)$$

$$V_{cur} = V_d \text{ if } V_d \leq V_{cur} < V_{d+1} \quad (20)$$

$\delta V$ and $\delta J$ were chosen to be sufficiently small to allow a good trade off between accuracy and runtime/memory usage.

### Virus phylogeny and antigenic distance

Each successful infection created a new virus derived from the infecting parent virus. In other words, the total number of distinct viruses was equal to the cumulative incidence. As each virus was a direct descendent of its parent, the new infecting virus shared the properties of the parent at the time of infection, $t_0$. Five key properties were stored for each virus:

1. The binding avidity at time of infection, $V_0$
2. The current binding avidity, $V$
3. The ID of the parent virus, $j$
4. The antigenic distance from parent, $\delta_{ij}$ where $i$ is the virus and $j$ is the parent virus
5. The ID of the host that the virus infected (for implementation purposes)

The full virus phylogeny can easily be reconstructed using property 3 by taking each extant virus; creating a link to its parent virus; then creating a link to its parent's parent etc. until the root virus is found. Similarly, the antigenic distance between any two viruses can be calculated using the following algorithm:

**Algorithm 1:** Algorithm to find antigenic distance between two viruses

```
distance = 0
A = number of parent viruses to root of virus i
B = number of parent viruses to root of virus j
Swap A and B such that A is the virus with fewest ancestors to root
while A < B do
    p = parent of i
    distance = distance + distance between i and p
    i = p
    A = number of parent viruses to root of i
end
if i == j then
    return distance
end
p_i = parent of i
p_j = parent of j
distance = distance + (distance between i and p_i) + (distance between j and p_j)
/* Note that here, i == NULL implies that i has no parent virus (ie. is
   the root virus)                                                    */
while p_i ≠ NULL and p_j ≠ NULL and p_i ≠ p_i do
    i = p_i
    j = p_j
    p_i = parent of i
    p_j = parent of j
    distance = distance + (distance between i and p_i) + (distance between j and p_j)
    if p_i == p_j == NULL then
        distance = Inf
    end
end
return distance
```
